# Transposon activity eliminates a crucial fungal secondary metabolite cluster while preserving pathogenicity

**DOI:** 10.1101/2025.05.14.653967

**Authors:** Jelmer Dijkstra, Anouk C. van Westerhoven, Yinping Li, Carolina Aguilera-Galvez, Giuliana Nakasato-Tagami, Xiaoqian Shi-Kunne, Desalegn W. Etalo, Gert H.J. Kema

## Abstract

**Summary:**

- Transposable elements can significantly influence the genome dynamics of clonal fungal plant pathogens. Tropical Race 4 (TR4) is a clonal lineage within the *Fusarium oxysporum* species complex that poses a substantial threat to global banana production. However, how transposable elements shape TR4’s genome and adaptation remains underexplored.
- We investigated the activity and impact of FoHeli1, a Helitron transposable element in the TR4 lineage, focusing on a Mozambican TR4 strain (M1) through a combination of genome analyses, metabolite profiling, and infection assays.
- FoHeli1 activity has been very recent within the TR4 lineage and is likely still ongoing. This has resulted in large structural variations in M1, including the loss of the conserved biosynthetic gene cluster required for the production of fusaric acid. We demonstrate that this deletion abolishes fusaric acid production and alters secondary metabolite profiles, but does not affect pathogenicity.
- Our results emphasize the significance of transposable elements, particularly FoHeli1, in reshaping the genetic and metabolic landscape of TR4 and challenge existing assumptions about the role of fusaric acid in pathogenicity.

## Introduction

Transposable elements (TEs) are powerful drivers of genome evolution, shaping genetic diversity and adaptation across the Tree of Life (Hayward & Gilbert, 2022). These mobile genetic elements can comprise substantial portions of the host genome (Wells & Feschotte, 2020) and influence critical processes, such as gene expression, splicing, deletion, recombination, and epigenetic regulation (Hirsch & Springer, 2017; Catlin & Josephs, 2022; Hayward & Gilbert, 2022). In plant-pathogenic fungi, TEs often associate closely with virulence-related genes and genomic compartments (Ma *et al*., 2010; Rouxel *et al*., 2011; Faino *et al*., 2016; Bertazzoni *et al*., 2018), playing a key role in adaptation and virulence evolution (Seidl & Thomma, 2017; Fouché *et al*., 2022). This association is thought to facilitate fast adaptation in the arms race between pathogenic fungi and their hosts, particularly in clonal populations, by modulating gene stability and expression (Chuma *et al*., 2011; de Jonge *et al*., 2013; Möller & Stukenbrock, 2017; Fouché *et al*., 2020; Torres *et al*., 2020; Fouché *et al*., 2022). However, while beneficial for short-term adaptation, the close association may have long-term negative fitness consequences due to TE accumulation, subsequent genome instability, and deleterious TE insertions (Fouché *et al*., 2022).

The importance of TEs in fungal genome evolution is particularly evident in the *Fusarium oxysporum* species complex (FOSC). FOSC members can infect over 100 plant hosts (Dean *et al*., 2012) and possess TE-enriched genome compartments that contribute to their adaptability and virulence (Ma *et al*., 2010; Li *et al*., 2020; Ayukawa *et al*., 2021; Dijkstra *et al*., 2024; van Westerhoven *et al*., 2024). Among FOSC lineages, the clonal Tropical Race 4 (TR4) lineage poses a severe threat to global banana production, a crop crucial for food security and local economics (Frison, 2004; Ordóñez *et al*., 2015; Ploetz *et al*., 2015), particularly in regions such as the Great Lakes area of Eastern and Central Africa (Karamura, 1998; Karangwa *et al*., 2016; van Westerhoven *et al*., 2022). TR4 is pathogenic to most banana cultivars, including the dominant cultivar Cavendish (Ploetz, 2015), while Race 1 and Race 2 strains continue to threaten locally produced banana varieties (Karangwa *et al*., 2016; Martínez-de la Parte *et al*., 2024). Despite the available genomic data on TR4, the impact TEs have on its genome evolution and adaptability have seen very limited investigation.

Helitrons, a distinct class of DNA transposons, are widespread across eukaryotic lineages and replicate via a rolling circle or peel-and-paste mechanism (Thomas & Pritham, 2015; Wells & Feschotte, 2020; Li *et al*., 2024). They have been extensively studied in plants (Du *et al*., 2009; Hu *et al*., 2019; Barro-Trastoy & Köhler, 2024) and animals (Grabundzija *et al*., 2016; Barro-Trastoy & Köhler, 2024). Helitrons are also present in fungal genomes, but have seen less investigation compared to plants and animals (Castanera *et al*., 2014; Castanera *et al*., 2016; Chellapan *et al*., 2016; Li *et al*., 2024). Here, we address this gap by investigating the recent activity and impact of Helitron element FoHeli1 in the Mozambican TR4 strain M1, providing experimental evidence of large Helitron-driven structural variations.

We identified two major deletions on core chromosomes caused by FoHeli1 transposition, including a 464 kb deletion that eliminates the entire 12-gene biosynthetic gene cluster (BGC) for fusaric acid (FA) production (Niehaus *et al*., 2014; Brown *et al*., 2015; Studt *et al*., 2016). As the first phytotoxin isolated from infected plants (Gaumann, 1957), FA has been implicated in virulence against tomato, banana and even mice (López-Díaz *et al*., 2018; Liu *et al*., 2020). Moreover, FA plays a role in the rhizosphere microbiome modulation in tomato (Jin *et al*., 2024). However, its role in virulence appears context-dependent, as FA production is dispensable for *Fusarium verticillioides* during maize infection (Brown *et al*., 2015).

Our study highlights the impact of Helitron activity on TR4 genome evolution, demonstrating their potential to reshape the genetic and metabolic landscape. We characterize FoHeli1 dynamics in the M1 strain and the reference TR4 strain II5, revealing extensive copy number variation across banana-infecting *Fusarium* strains. Furthermore, we assess the impact of the FA BGC loss on growth, metabolism, and pathogenicity, underscoring the broader implications of TE-mediated genomic changes on fungal biology and host-pathogen interactions.

## Materials and methods

### Fungal strains, growth conditions and transformation

*Fusarium* Tropical Race 4 reference strain II5, strains M1 and M2 isolated from Mozambique (van Westerhoven *et al*., 2023; van Westerhoven *et al*., 2024) and newly generated *fub1* knockout strain *Δfub1* and *fub1^c^* complementation mutants were used in this study. Strains were stored as conidial suspension stocks in 15% glycerol at -80 °C. General culturing of the fungal strains was performed using potato dextrose agar (PDA) at 25 °C.

The *Δfub1* mutant was generated in the II5 background as previously described (Dijkstra *et al*., 2024). Complementation was performed using the same procedure instead using a DNA fragment containing the *fub1* gene and 1kb of upstream and downstream DNA fused with a hygromycin cassette by overlap extension PCR as donor fragment. Complementation was done by ectopic integration using the same procedure as in Dijkstra et al., (2024) in the absence of a Cas9 vector. Used primers are listed in Table S1.

### Whole genome comparison

The previously published whole genome sequences of strains II5 and M1 were used for analysis (van Westerhoven *et al*., 2024). Putative biosynthetic gene clusters were predicted for the genomes of strains II5 and M1 using Antismash 7.1 with standard relaxed detection strictness (Blin *et al*., 2023). The FA BGC cluster and Starship representation were made using Clinker (Gilchrist & Chooi, 2021). Chromosome alignments were made using syntenyPlotteR (Quigley *et al*., 2023).

### Transposable element analysis

Initial FoHeli1 identification was done using BLAST (Camacho *et al*., 2009) and comparison to known FoHeli1 sequences (Chellapan *et al*., 2016). TEs in long-read sequenced strains II5 and M1 were annotated and analyzed using Earl Grey using the standard parameters (Baril *et al*., 2024). Full-length FoHeli1 copies were aligned using Clustal Omega to determine the number of SNPs between each copy (Sievers *et al*., 2011). The FoHeli1 chromosomal location plot was made with the use of MG2C (Chao *et al*., 2021). FoHeli1 copy numbers were further analyzed in 58 short-read sequenced *Fusarium* strains isolated from banana using DeviaTE (Weilguny & Kofler, 2019). The sequence of II5 translation elongation factor 1-alpha (EF-1 alpha) was used as single copy reference gene for copy number estimation. All used strains and their FoHeli1 copy number are noted in Table S2.

DNA from strains II5, M1, 36102, and CR1.1 was extracted from freeze-dried mycelium using the Masterpure™ Yeast DNA Purification Kit (LGC Biosearch™ Technologies, Hoddesdon, United Kingdom). Genomic DNA was used for PCR and subsequent gel electrophoresis to check for FoHeli1 circular intermediates. The primers used are listed in Table S1.

### Exometabolite extraction

Strains II5, M1 and M2 were grown on PDA plates for 7 days. Five agar plugs were then used to inoculate flasks containing 100 ml PDB supplemented with 5% banana root exudate. Cultures were grown in triplicate at 25 °C and 150 rpm for 7 days. PDB with 5% banana root exudate without *Fusarium* was used as control. After growth the culture filtrates were obtained by filtering the medium through a 0.2 µm filter. Culture filtrates were freeze dried and afterwards dissolved in 4 mL ethyl acetate for one hour on a rolling shaker. Samples were subsequently dried with nitrogen and dissolved in 5 mL 75% MeOH, 0.1% formic acid. Afterwards samples were sonicated for one hour. Samples were further centrifuged twice at 13.000 rpm for 10 minutes. HPLC vials were filled with 250 µL supernatant for chromatographic analysis.

### Exometabolite analysis

An UltiMate 3000 U-HPLC system (Dionex) was used to create a 45-minute linear gradient of 5-35% (v/v) acetonitrile in 0.1% (v/v) formic acid (FA) in water at a flow rate of 0.19 mL min^−1^. 5 µL of each extract was injected, and compounds were separated on a Luna C18 column (2.0 x 150 mm, 3µm; Phenomenex, Torrance, California USA) maintained at 40 °C (De Vos *et al*., 2007). The detection of compounds eluting from the column was performed with a Q-Exactive Plus Orbitrap FTMS mass spectrometer (Thermo Scientific™, Waltham, Massachusetts USA). Full scan MS data were generated with electrospray in switching positive/negative ionisation mode at a mass resolution of 35,000 (FWHM at m/z 200) in a range of m/z 95-1350. Subsequent untargeted MS/MS analyses were performed with separate positive or negative electrospray ionisation at a normalised collision energy of 27 and a mass resolution of 17,500. The ionisation voltage was optimised at 3.5 kV for positive mode and 2.5 kV for negative mode; the capillary temperature was set at 250 °C; the auxiliary gas heater temperature was set to 220 °C; sheath gas, auxiliary gas, and the sweep gas flow were optimised at 36, 10 and 1 arbitrary units, respectively. Automatic gain control was set at 3e^6^ and the injection time at 100 ms. External mass calibration with formic acid clusters was performed in both positive and negative ionisation modes before each sample series.

#### LCMS data preprocessing and analysis

Mass peak picking and alignment were performed using Metalign software (Lommen, 2009). Mass features in the resulting peak list were considered as a real signal if they were detected with an intensity of more than 3 times the noise value and in 2 out of the three biological replicates of at least one treatment. Mass features originating from the same metabolites were subsequently reconstituted based on their similar retention window and their intensity correlation across all measured samples, using MSClust software (Tikunov *et al*., 2012). This resulted in the relative intensity of 725 putative mass features potentially representing individual metabolites across the samples. The raw data were processed using R (4.4.1 [2024-06-14]) for gap-filling and normalisation. Average PDB intensities were calculated for each feature, and these were subtracted from the sample and QC intensities to correct for baseline signals. Negative values were replaced with small positive random values between 90 and 100 to avoid computational errors during subsequent analysis. Logarithmic transformation (log2) was applied to normalised intensities to stabilise variance and normalise distributions. Differentially abundant metabolites identified through statistical analyses were subjected to structural annotation. Representative samples from each treatment were analysed using untargeted positive and negative ionisation MS/MS, generating fragmentation spectra that were processed and exported as .mgf files in MZmine 4.3.0 (Schmid *et al*., 2023). These spectra were processed using SIRIUS software for molecular formula prediction and compound annotation based on fragmentation patterns (Dührkop *et al*., 2019). MS/MS spectra were matched against public databases such as Biocyc, COCONUT, KEGG, KNApSAcK, LOTUS, GNPS, HMDB, and PubChem:bio and metabolites to enhance confidence in annotation. Annotation confidence levels were categorised following the Metabolomics Standards Initiative (MSI) guidelines, with Level 1 corresponding to confirmed identifications through reference standards (only for Fusaric acid) and Levels 2-4 representing varying degrees of confidence based on spectral similarity and predictive algorithms.

#### Principal Component Analysis (PCA)

Principal Component Analysis was conducted to visualise variance among the experimental groups. The normalised intensity data were transposed, and PCA was performed using the prcomp function in R with data scaling. The PCA results were visualised using fviz_pca_ind from the factoextra package. Two PCA analyses were performed, including all experimental groups and excluding PDB samples, to assess group-specific variation.

#### Differential Analysis

Differential metabolite abundance between experimental groups (M1 vs. M2, M1 vs. II5, and M2 vs. II5) was evaluated using the limma package in R. For each comparison, a linear model was fitted to the data, and empirical Bayes moderation was applied to improve robustness. The adjusted p-value threshold (false discovery rate, FDR) was set at 0.05. Fold changes greater than ±2 were considered biologically significant. All annotated features in the comparison between II5 and M1 are noted in Table S3. Annotated features in the comparison between II5 and M2 are noted in Table S4.

#### Volcano Plot Visualization

Volcano plots were generated to highlight significant metabolites for each pairwise comparison. Log fold changes (logFC) and -log10 transformed adjusted p-values were plotted using ggplot2. Significant features were colour-coded by pathway categories (e.g., alkaloids, carbohydrates, terpenoids) with a custom colour palette. Threshold lines at logFC ±1 and p-value < 0.05 were annotated to demarcate significance criteria.

### Growth assays

For the determination of colony growth, agar plugs were taken from glycerol stock inoculated PDA plates and put on fresh PDA, CDA or PDA plates supplemented with 200 ppm fusaric acid (Sigma-Aldrich, Saint Louis, USA). Colony diameter was measured at 3 days post inoculation (dpi). Colonies were photographed at 3 dpi. Photos for colony pigmentation were taken at 8 dpi.

### Plant growth conditions and infection assays

Cavendish plants were inoculated as described previously (García-Bastidas *et al*., 2019). In brief, conidia were produced by inoculating flasks of Mung bean medium with agar plugs of the different wild-type *Fusarium* strains and mutants. The flasks were incubated for 5 days at 150 rpm 25 °C. Conidia were collected by filtering the medium through two layers of sterile cheesecloth. The conidia were subsequently quantified and diluted to the desired concentration. Prior to inoculation, Cavendish plants were wounded in their roots twice and inoculated with 200 mL of either 1·10^6^ or 1·10^4^ conidia/ml by pouring. Plants were monitored and scored after 8-10 weeks post-inoculation. To assess virulence levels, plants were cut longitudinally at the corm, and pictures were taken. Finally, the percentage of necrosed corm tissue was determined using ImageJ.

## Results

### Helitron activity caused loss of the fusaric acid biosynthetic gene cluster in M1

TR4 has recently been shown to be spreading beyond its initial incursion point in Mozambique (van Westerhoven *et al*., 2023). The TR4 strains from Mozambique were generally found to be highly similar to other globally occurring TR4 strains (van Westerhoven *et al*., 2023). However, whole genome comparisons with the TR4 reference genome reveals two large scale deletions for strain M1, which was previously used for long-read whole genome sequencing (van Westerhoven *et al*., 2023; van Westerhoven *et al*., 2024). These two large-scale deletions encompass 184 and 464 kb of the sub-telomeric regions of the M1 core chromosomes 2 and 3, respectively (Fig. 1a), representing approximately 1.25% of the 51 Mb sized genome of II5 (Dijkstra *et al*., 2024). Together, the deleted regions comprise 210 putative genes, showing a significant loss of genetic material.

**Fig. 1.**
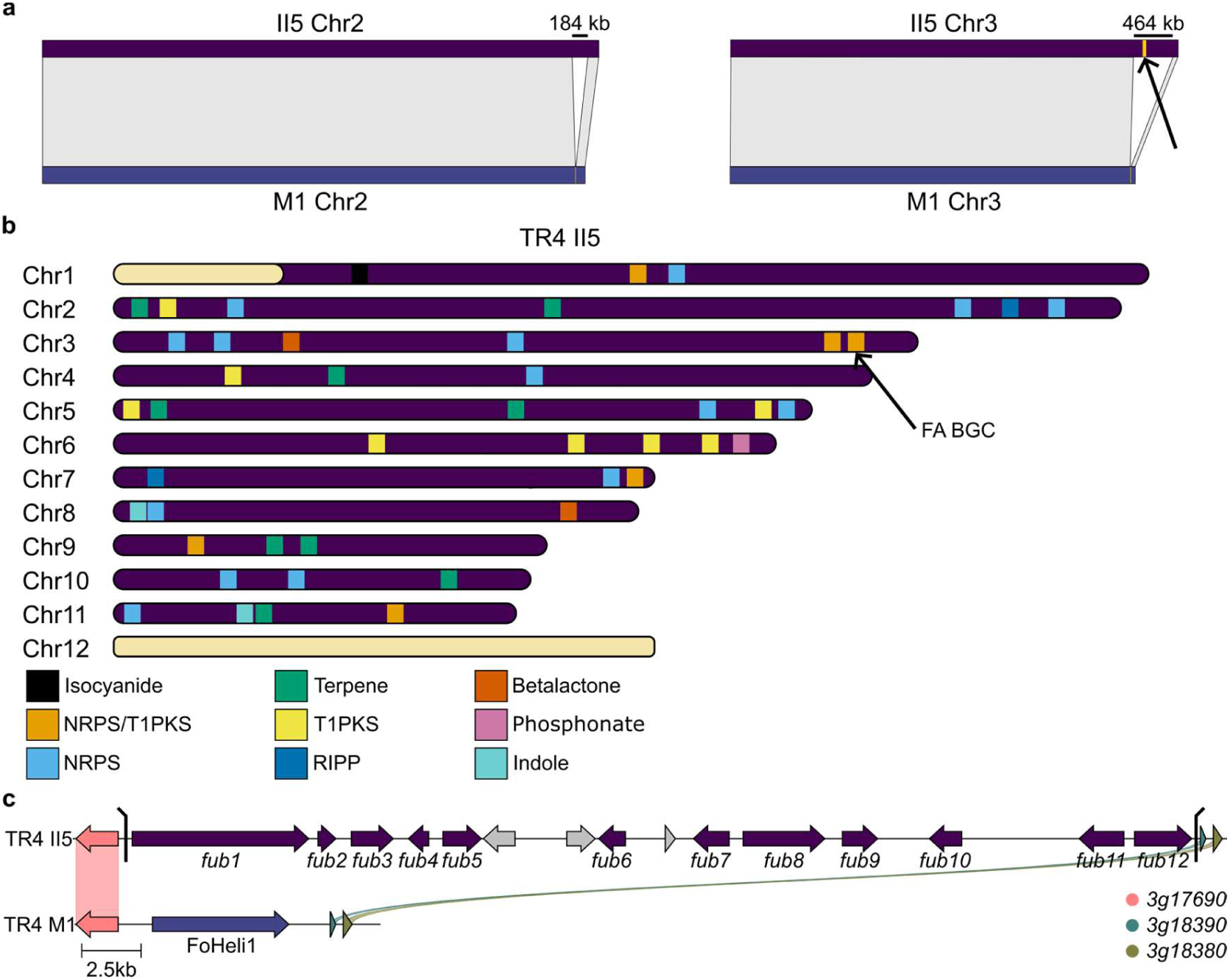
Two large-scale deletions including the FA BGC on core chromosomes in TR4 strain M1. **a**) Alignment of core chromosomes 2 and 3 between TR4 strains II5 and M1. The approximate length of the deleted sequences compared to II5 is shown. The approximate location of the fusaric acid biosynthetic gene cluster (FA-BGC) is marked yellow on II5 chromosome 3 (black arrow). **b**) Schematic representation of the approximate locations of all predicted BGCs (46) and their types in TR4 strain II5. NRPS=Non ribosomal peptide synthase, T1PKS=Type I polyketide synthase, RIPP=Ribosomally synthesized and post-translationally modified peptide. The FA-BGC is located at the end of chromosome 3 (black arrow). **c**) Alignment of the FA-BGC regions in TR4 strains II5 and M1. For TR4 strain II5, the FA-BGC and its 12 genes are depicted, as well as the genes flanking the deletion. Black lines indicate the rest of the deleted sequence compared to II5. The entire FA-BGC of TR4 strain M1 is replaced by the ∼6kb FoHeli1 TE.

Prediction of BGCs in the reference genome of II5 indicates that the entire FA BGC is located within the borders of the deletion on chromosome 3 (Fig. 1b). Further genomic analysis, BGC prediction, and PCR confirmed that M1 had lost the 12 *fub* genes comprising the FA BGC and that the cluster was not translocated elsewhere in the genome (Fig. S1). To determine the potential cause of these large-scale deletions, we examined the deletion sites on chromosomes 2 and 3 and found that both deleted sequences were replaced by a short sequence of approximately 6 kb. Further BLAST analysis of this fragment indicated the presence of a FoHeli1 Helitron, a TE that replicates by a rolling-circle mechanism (Schmidt *et al*., 2013; Chellapan *et al*., 2016; Barro-Trastoy & Köhler, 2024). The FoHeli1 copy on chromosome 3 shares 98.6% identity with FoHeli1 in the tomato infecting strain *Fol*4287 (MF805714.1). We conclude that a total of 648 kb was lost in M1 because of FoHeli1 activity, including the full-length FA BGC (Fig. 1c).

To check whether loss of the FA BGC has occurred in other strains, we performed a blast analysis of the key biosynthesis gene *fub1* across a set of 35 publicly available *Fusarium oxysporum* genome assemblies (Meng *et al*., 2024). This analysis shows the loss of the FA BGC in four additional strains that are pathogenic to common bean (Meng *et al*., 2024). However, for these strains, no traces of FoHeli1 are present in the deleted region. Instead, a large 174 kb region is observed that is absent in II5 (Fig. S2a). Based on the large size of this fragment, in the otherwise conserved core chromosomes, we further analyzed the genes present in this region for strain F109. Based on the presence of several key genes, we conclude that this large section that is absent in II5 encompasses a Starship element (Fig.S2a-b) (Gluck-Thaler *et al*., 2022). These results indicate that the FA BGC might be lost more commonly through different processes.

### FoHeli1 activity in the TR4 lineage is recent and ongoing

The observed deletions indicate that Helitrons play an important role in the genome dynamics of *Fusarium oxysporum*. To further characterize FoHeli1 dynamics and search for additional copies, all TEs in the chromosomal assemblies of isolates II5 and M1 were predicted using Earl Grey (Baril *et al*., 2024) (Fig. S3), which detected 15 and 16 copies of FoHeli1 TEs in II5 and M1, respectively. Most copies are full length (∼6 kb), but both isolates contain a single copy of very short length corresponding to approximately the first 230 bases of the full-length FoHeli1 copies. This short length sequence might represent the remnants of a (non-autonomous) FoHeli1. Similarly, II5 and M1 contain one and two copies of FoHeli1, respectively, with a 5’ truncation of several hundred bases, which may prevent further transposition. Transposable elements tend to accumulate mutations over time. By calculating the Kimura distance between TE sequences within the same family, we obtained a relative indication of when these elements last actively transposed (Chalopin *et al*., 2015). FoHeli1 copies have the lowest average weighted Kimura distance in both II5 (0.04) and M1 (0.03) in comparison to all other predicted TEs which have a median average Kimura distance of 11.44 and 12.08 for II5 and M1, respectively, which suggests very recent activity (Fig. 2a – Fig. S4). To analyse the similarity of FoHeli1 copies in the strains, we compared the sequences of all individual copies. In general, the copies cluster at the strain level, except for two M1 derived copies that cluster along with the II5 derived copies (Fig. 2b). Interestingly, the FoHeli1 copies located on accessory chromosome 12 of II5 formed a distinct cluster, suggesting intrachromosomal transpositions (Fig. 2b). Moreover, we observe that the vast majority of the FoHeli1 copies were found on different chromosomal locations between II5 and M1, further indicating recent activity of this TE (Fig. 2c). For both isolates, accessory chromosome 12 contains the most FoHeli1 copies in line with their generally higher TE density. Another way to determine FoHeli1 activity is to search for copies with multiple 5’ termini, which can arise from self-insertion events (Chellapan *et al*., 2016). We used the previously determined FoHeli1 5’ terminal sequence (Chellapan *et al*., 2016) and found that, of the 14 longer copies in II5, one copy contains two 5’ termini while another copy contains five 5’ termini. In M1, on the other hand, 5 out of 15 of the longer FoHeli1 copies contain two 5’ termini, while three copies contain three 5’ termini (Fig. S5). PCR further confirmed the presence of a circular intermediate of FoHeli1 in both TR4 isolates, while being absent in the other banana infecting *Fusarium* strains 36102 and CR1.1, which do not belong to the clonal TR4 lineage and lack any FoHeli1 copies (Fig. 2d). We also tested how recent FoHeli1 activity took place as well as its propensity to be involved in genomic changes in the TR4 lineage in strains UKTR4 and HN51, where FoHeli1 also had the lowest average weighted Kimura distance and was, in several cases, located at the border of rearrangements (Fig. S6), one of which was independently observed previously (Zhang *et al*., 2024). Taken together, we show that FoHeli1 activity in the TR4 lineage is very recent and most likely still ongoing.

**Fig. 2.**
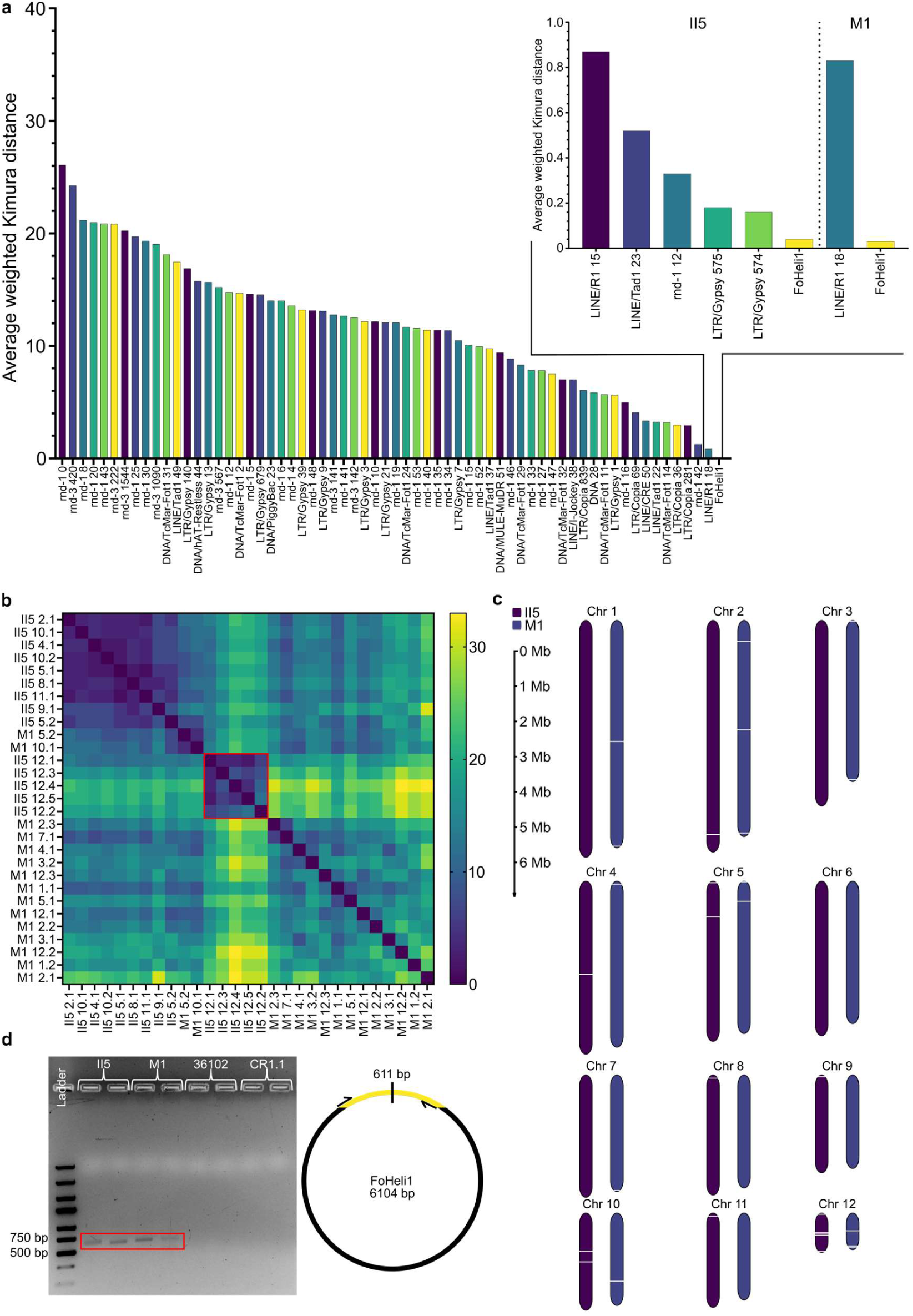
Recent FoHeli1 activity in the TR4 lineage. **a**) Bar chart showing the average weighted Kimura distance of all TEs predicted and calculated by Earl Grey in TR4 strain M1. Inset shows TEs with average weighted Kimura distance ≤1 in TR4 strains II5 and M1. **b**) Heatmap showing the number of SNPs between all long FoHeli1 copies in TR4 strains II5 and M1. **c**) Schematic representation of the location of all FoHeli1 copies per chromosome of TR4 strains II5 and M1. FoHeli1 locations are marked by white gaps. **d**) Gel electrophoresis of a PCR using FoHeli1 primers for a circular template (depicted right) on TR4 strains II5, M1, 36102 and Race 1 strain CR1.1. The expected band size is 611 bp.

### FoHeli1 copy number is highly diverse in banana-infecting *Fusarium* strains

We used DeviaTE (Weilguny & Kofler, 2019) to estimate the FoHeli1 copy number of 58 Illumina sequenced banana infecting *Fusarium* strains (Martínez-de la Parte *et al*., 2024; van Westerhoven *et al*., 2024). These estimates show that the copy number of FoHeli1 ranges between 9 and 36 in TR4 strains (Fig. 3a). Interestingly, the highest copy numbers (N=25-36) are observed in a set of TR4 strains which have intrachromosomal duplications of accessory chromosome 12 (Fig. 3a) (Dijkstra *et al*., 2024). The copy number estimates for isolate II5 by DeviaTE are higher (N=25) than the copy number obtained from Earl Grey (N=15). This is caused by intrachromsomal duplications of chromosome 12, that is not resolved in the assembly (Dijkstra *et al*., 2024). In contrast to TR4, Race 1 and Race 2 strains show a much wider range of FoHeli1 copy number than TR4 (Fig. 3b). Some of the analyzed Race 1 strains do not have any FoHeli1 copies, whereas all Race 2 strains contain a high number of FoHeli1 copies (N>50). It has previously been observed that Race 2 strains have a larger genome size and contain more accessory material (van Westerhoven *et al*., 2024). Comparing the genome size of 47 banana-infecting *Fusarium* strains with their FoHeli1 copy number shows a moderate positive correlation between the two factors (Fig. S7). This indicates that the larger number of FoHeli1 elements in Race 2 strains is possibly caused by a larger accessory genome size (van Westerhoven *et al*., 2024). Helitrons are known to occasionally capture genes or gene fragments during transposition (Castanera *et al*., 2014; Barro-Trastoy & Köhler, 2024). Hence, we searched for such events but identified only a single case in Race 2 strain C081 (Fig. 3c). However, such gene capture events are likely to be missed in Illumina sequenced strains due to the limitations of short-read sequencing for structural genomic analyses. Interestingly, FoHeli1 copies are present in all TR4 strains, which all contain chromosome 12, but FoHeli1 is absent in the most closely related strain 36102, which lacks this chromosome (van Westerhoven *et al*., 2024), thus raising questions on the origin of FoHeli1 within the TR4 lineage.

**Fig. 3.**
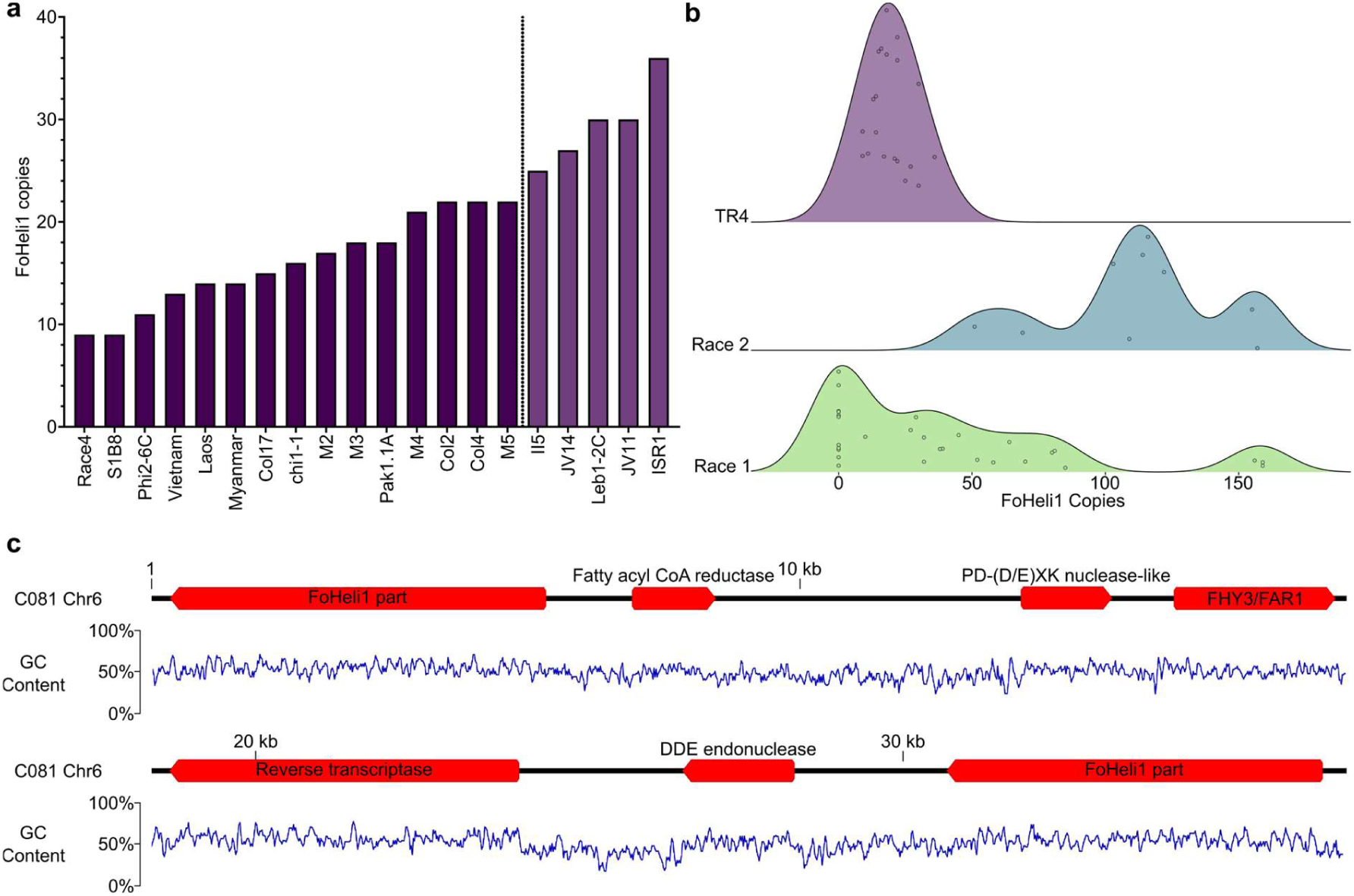
FoHeli1 copy number is diverse in banana infecting *Fusarium* strains. **a**) Estimated FoHeli1 copy number in 20 Illumina sequenced TR4 strains based on DeviaTE. Strains with duplications within accessory chromosome 12 (Dijkstra *et al*., 2024) are noted with a lighter shade and separated by a dotted line. **b**) Ridgeline plot of estimated FoHeli1 copy number in 58 Illumina sequenced banana infecting *Fusarium* strains based on DeviaTE. The plot separates Tropical Race 4 (TR4), Race 1 and Race 2 strains. Each dot represents one strain. The estimated copy number was cut off to the nearest full number for each strain. **c**) Fragment of the genomic sequence of Race 2 strain C081 (∼38 kb) depicting a gene capture event by a FoHeli1 copy. Domains of the captured genes are noted. GC content is plotted below.

### LC-MS analysis of M1 confirms the absence of fusaric acid

To investigate how the large-scale deletions on chromosomes 2 and 3 affect the overall metabolome, particularly FA biosynthesis, we performed untargeted metabolomics profiling on exometabolites of strains II5 and M1 following *in vitro* cultivation. Firstly, comparison to a commercial FA standard confirmed the presence of FA and the derivative dehydrofusaric acid in II5 and their absence in M1 (Fig. 4a). Thus, the deletion because of FoHeli1 activity abolished FA production in M1.

**Fig. 4.**
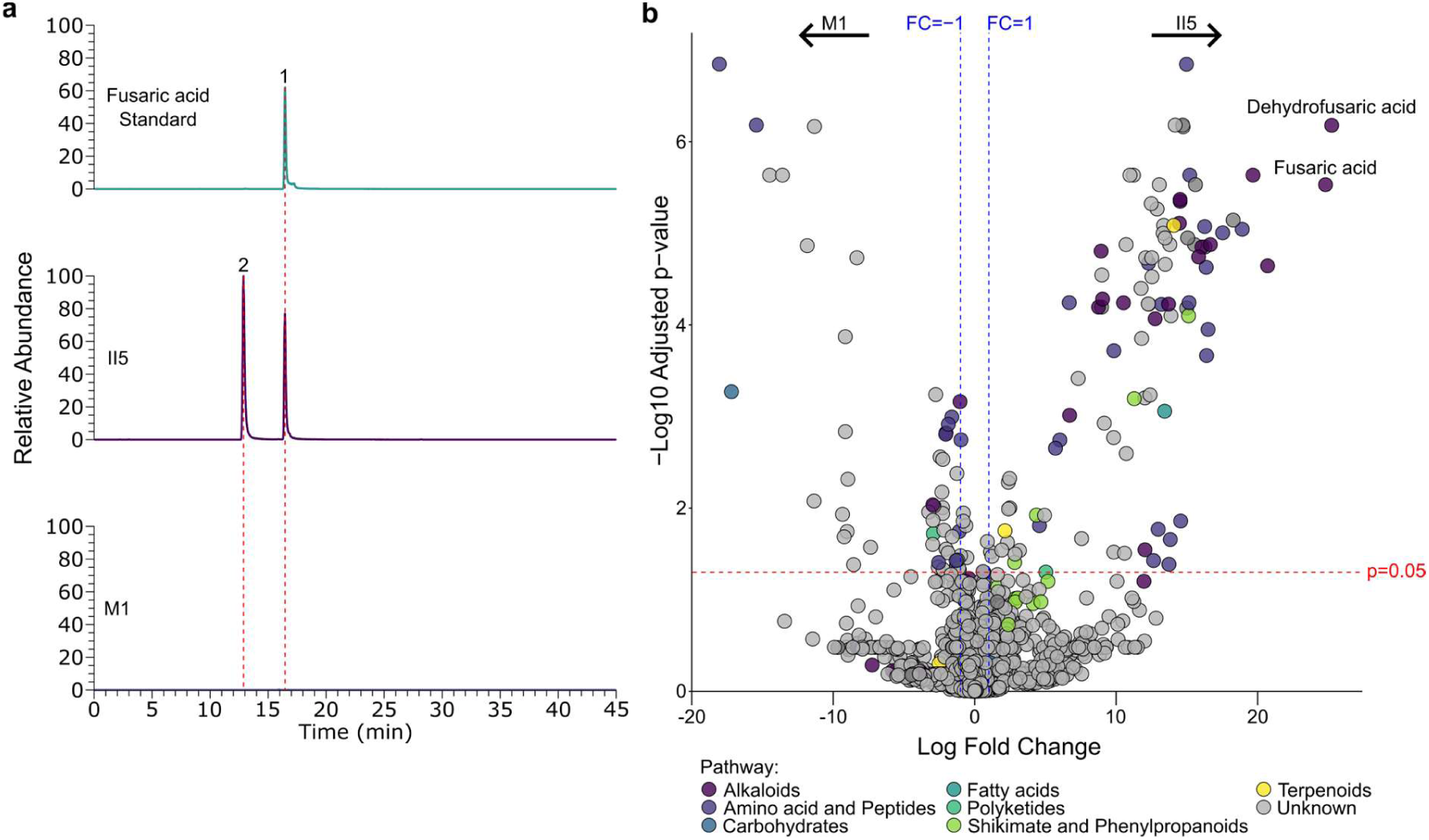
TR4 strain M1 does not produce fusaric acid. **a**) LC-MS profile of commercial fusaric acid standard and liquid cultures of TR4 strains II5 and M1. Peak 1 corresponds to fusaric acid, and peak 2 corresponds to dehydrofusaric acid (red dotted lines). **b**) Volcano plot of metabolite features detected in TR4 strains II5 and M1. Blue threshold lines are shown at a log fold change (FC)>1 and FC<-1. The red threshold line indicates an adjusted p-value of 0.05. Negative fold change features are more abundant in M1; positive fold change features are more abundant in II5.

In total, 1594 metabolite features were detected in positive and negative ionization modes across the samples, including the growing medium (PDB) (Fig. 4b). We performed Principal Component Analysis (PCA) to examine the variation in the exometabolite profile among strains II5, M1 and M2 and the growing medium. The first and the second principal components explained 46.2% and 15.3% of the variability, respectively, accounting for a cumulative 61.5% of the total variability in the metabolite data (Fig. S8). The first principal component underscores the differences in metabolites between the growing medium and the fungal isolates, reflecting the metabolic activity of the fungi. In contrast, the second principal component depicts the variation in metabolite profiles between the three TR4 isolates. Additionally, a pairwise comparison of the individual metabolite features between II5 and M1 revealed that 146 features exhibited significant variation (FC>1/FC<-1 and P <0.05), which included compounds from the alkaloids, amino acid and peptides, carbohydrates, fatty acids, polyketides, shikimate and phenylpropanoids, terpenoid pathways (Fig. 4b). In contrast, pairwise comparison between II5 and M2 showed 107 metabolite features with significant variation. M2 did not show significantly different FA production levels. In particular, comparative metabolomics of M1 with II5 showed either a complete absence or a significant reduction in metabolites belonging to the alkaloid pathway, including fusaric acid, dehydrofusaric acid, and other derivatives (Fig. 4b). Moreover, several metabolites associated with dipeptides showed a significantly higher abundance in II5.

### Loss of the fusaric acid biosynthetic gene cluster impacts fungal development

FA has been shown previously to have an antimicrobial effect, including self-inhibition of the producing fungus (Crutcher *et al*., 2015; Studt *et al*., 2016). Normally, fungal self-protection against FA occurs by the cluster-specific transporter (*fub11*) and the regulation of derivatives with reduced toxicity (*fub12*) (Studt *et al*., 2016). As M1 has lost the entire FA BGC, both functions should be disrupted. In turn, we observed an elevated sensitivity to exogenously applied FA in M1 compared to II5 (Fig. 5a-b). Similarly, FA production has been previously implicated with growth and biomass levels (Crutcher *et al*., 2015; Studt *et al*., 2016). To check for potential growth effects, we performed growth experiments using II5, M1, the *fub1* knock-out strain Δ*fub1*, and the complementation strain Δ*fub1*^c^. Growth on regular PDA or FA production promoting Czapek-Dox agar (CDA) showed no significant differences between all tested strains. Only M1 showed slightly higher growth rates on PDA plates similar to the experiment using exogenous FA application (Fig. 5c). Loss of FA production has been shown previously to affect the production of other secondary metabolites such as the pigment bikaverin (López-Díaz *et al*., 2018; Phasha *et al*., 2021). Similarly, we also observe more pigmentation of the M1 and Δ*fub1* colonies compared to II5 and Δ*fub1*^c^ (Fig. 5d).

**Fig. 5.**
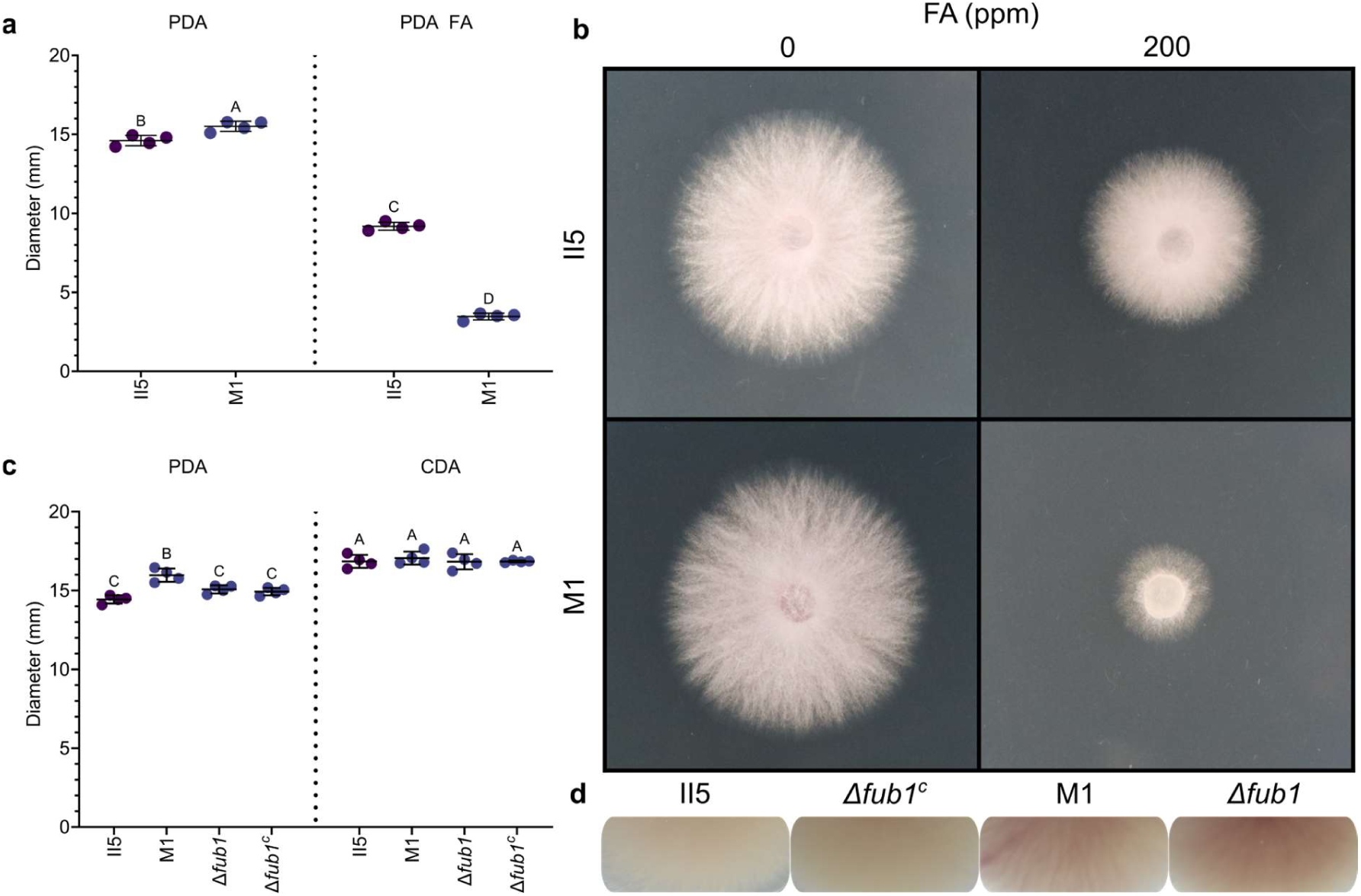
Fusaric acid biosynthetic gene cluster loss affects different phenotypes. **a**) Colony diameter (mm) of TR4 strains II5 and M1 on PDA and PDA plates supplemented with 200 ppm fusaric acid. Growth was quantified at 3 days post inoculation (dpi). Letters indicate significant differences between treatments (Tukey-Kramer test; P<0.05). Data are shown as mean ±SD (n=4). **b**) PDA plates and PDA plates supplemented with 200 ppm fusaric acid inoculated with TR4 strains II5 or M1 photographed at 3 dpi. **c**) Colony diameter (mm) of PDA plates and CDA plates inoculated with TR4 strains II5, M1, Δ*fub1* mutant and complementation. Growth was quantified at 3 dpi. Letters indicate significant differences between treatments (Tukey-Kramer test; P<0.05). Data are shown as mean ±SD (n=4). **d**) Photos of the colony edge for strains II5, Δ*fub1^c^,* M1 and Δ*fub1* inoculated on PDA plates at 8 dpi. Photographed from the bottom of the plate.

### Fusaric acid is not required for pathogenicity on banana

To test whether the large deletions affect M1 virulence and if FA contributes to banana infection, Cavendish plants were inoculated with wildtype strains II5 and M1, as well as the newly generated Δ*fub1* mutant and the complemented strain Δ*fub1*^c^. The results from the infection assays showed that isolates II5 and M1 caused similar disease severity irrespective of the used spore concentration (Fig. 6a). Similar results were obtained with the FA producing M2 strain (Fig. S9). Comparable to the M1 infection assay, no significant difference was observed between II5, the Δ*fub1* mutant, and the Δ*fub1^c^* complementation line at the lower spore concentration (Fig. 6b). In general, a lower spore concentration leads to lower corm necrosis levels and reduced or delayed disease incidence and does not lead to significant differences between the different strains and mutants. No significant difference was observed between II5 and the Δ*fub1* mutant at the regular spore concentration (10^6^) either (Fig. S10). Together, these results indicate that the loss of the major toxin FA by FoHeli1 activity or CRISPR/Cas-9 mediated knockout of core enzyme *fub1* does not affect the overall virulence of TR4 on Cavendish plants.

**Fig. 6.**
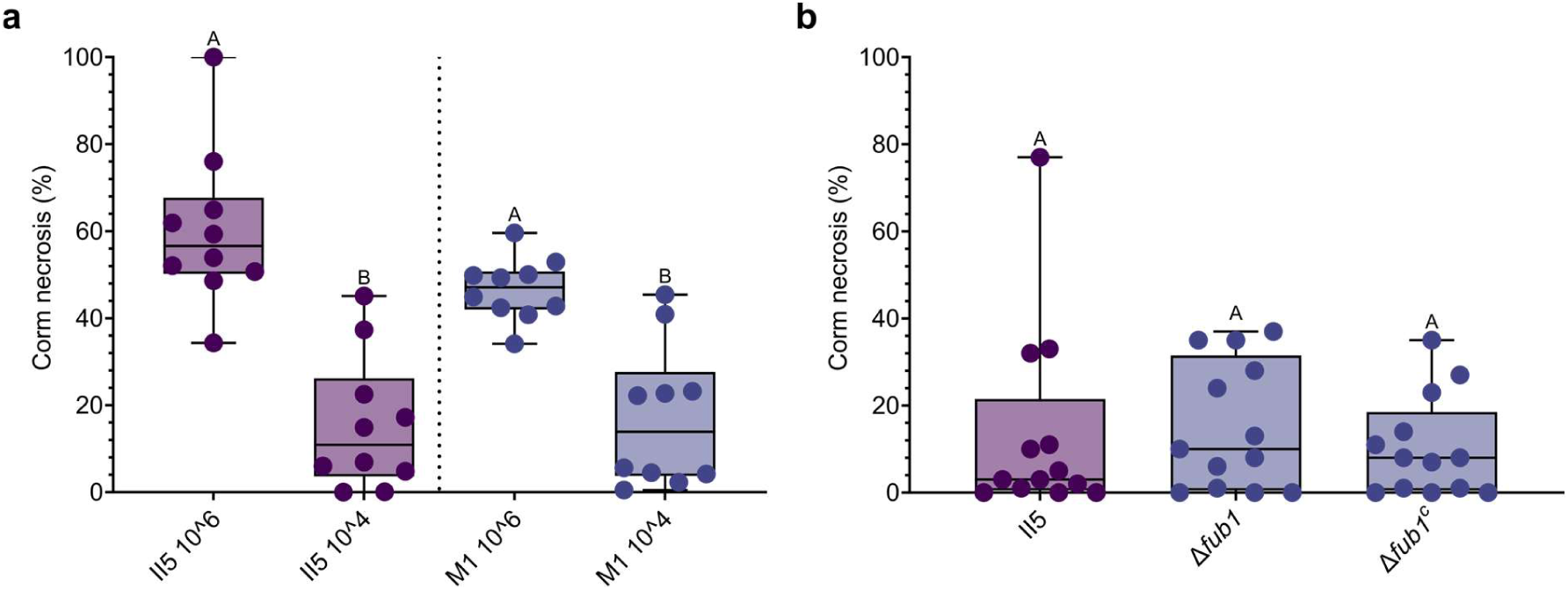
Fusaric acid is not required for virulence on banana. **a**) Percentage of corm necrosis at 10 weeks post-inoculation of Cavendish ‘Grand Naine’ plants inoculated with the reference TR4 strain II5 or strain M1. Plants were inoculated with either 10^6^ or 10^4^ spores per mL. Corm necrosis was quantified using ImageJ (n=10). Letters indicate significant differences between treatments (Tukey-Kramer test; P<0.05). **b**) Percentage of corm necrosis at 10 weeks post-inoculation of Cavendish ‘Grand Naine’ plants inoculated with strain II5, Δ*fub1* mutant or the Δ*fub1^c^* complementation strain. All plants were inoculated with 10^4^ spores per mL. Corm necrosis was quantified using ImageJ (n=13). Letters indicate significant differences between treatments (Tukey-Kramer test; P<0.05).

## Discussion

Our study reveals the genomic and metabolic dynamics driven by Helitron activity in the banana infecting clonal TR4 lineage. By characterizing large-scale deletions mediated by the transposon FoHeli1 through bioinformatic analysis, metabolomic profiling, growth and pathogenicity assays, our work underscores the significant impact of transposable elements on genome dynamics and the adaptive potential of fungal pathogens. These deletions, notably the loss of the fusaric acid biosynthetic gene cluster, provide new insight into understanding the evolutionary trade-offs inherent in TE-driven genome remodeling.

Helitrons have been implicated to play a role in genomic variation across diverse eukaryotes (Wells & Feschotte, 2020; Hayward & Gilbert, 2022; Barro-Trastoy & Köhler, 2024; Li *et al*., 2024). Although Helitrons have seen some investigation within fungi (Castanera *et al*., 2014; Li *et al*., 2024; Oggenfuss *et al*., 2024), including *Fusarium oxysporum* (Chellapan *et al*., 2016) their impact on genome dynamics in fungal pathogens has remained underexplored, particularly in banana-infecting *Fusarium*. The recent and likely ongoing activity of FoHeli1 underscores how transposable elements can generate genetic variation in clonally reproducing pathogens. The low sequence diversity among FoHeli1 copies in M1 and II5 strains, their occurrence in distinct loci between the two strains, and the detection of a putative circular intermediate suggest an active phase of transposition that continues to reshape the TR4 genome. This activity aligns with the prior observation of Helitron-mediated genomic changes in other *Fusarium* strains, such as the loss of an effector gene associated with host specificity (Biju *et al*., 2017), further highlighting the evolutionary significance of these elements in host-pathogen interactions. Another indication of ongoing FoHeli1 activity within the TR4 lineage is our observation of multiple 5’ termini in many M1 FoHeli1 copies, which are likely to arise following self-insertion events (Chellapan *et al*., 2016). The presence of FoHeli1 copies at the border of genome rearrangements in additional TR4 strains suggest a broader TR4 wide impact of FoHeli1 activity. However, some investigated strains lack FoHeli1 entirely, such as the less virulent strain 36102, this strain is most closely related to the clonal TR4 lineage and lacks accessory chromosome 12 (van Westerhoven *et al*., 2024). This raises important questions around a potential horizontal transfer origin for FoHeli1 within the TR4 lineage.

Although Helitrons have also been implicated in the capture and transposition of adjacent genes (Castanera *et al*., 2014; Barro-Trastoy & Köhler, 2024), we did not observe this within either II5 or M1. Putative captured genes, however, were observed for Race 2 strain C081. The availability of additional long-read genome assemblies could provide deeper insights into the frequency and mechanisms of gene capture by FoHeli1 and other Helitrons in banana-infecting *Fusarium*. Furthermore, such assemblies could improve understanding of FoHeli1-related structural genomic changes across the *Fusarium oxysporum* species complex (FOSC) and their evolutionary implications for host-pathogen interactions.

Further estimation of FoHeli1 copy number in 58 sequenced banana-infecting *Fusarium* strains indicates a large variation in copy number. TR4 strains recently shown to harbor intrachromosomal duplications of accessory chromosome 12 possess the highest number of FoHeli1 copies within the TR4 lineage (Dijkstra *et al*., 2024). Interestingly, all FoHeli1 copies on chromosome 12 of II5 are highly similar to one another and distinct from those on core chromosomes. These findings indicate a recent local transposition burst, as these copies are not part of any larger segmental duplications. Compared to TR4, Race 1 and Race 2 strains show a much wider range of FoHeli1 copy number variation. Although generally thought to replicate via a peel-and-paste mechanism (Barro-Trastoy & Köhler, 2024), experimental evidence in maize suggests transposition by excision can occur as well (Li & Dooner, 2009). With the occurrence of FoHeli1 in distinct loci in all investigated TR4 genome assemblies, the degree of copy number variation is more suggestive of predominant transposition by excision rather than a peel-and-paste mechanism.

The loss of the FA BGC challenges the prevailing view of FA as a critical virulence factor in the *Fusarium*-banana interaction (Li *et al*., 2013; Liu *et al*., 2020). While FA has been associated with phytotoxicity, self-toxicity, antimicrobial activity, and rhizosphere modulation (Bacon *et al*., 2006; Son *et al*., 2008; Crutcher *et al*., 2015; Bohni *et al*., 2016; Studt *et al*., 2016; López-Díaz *et al*., 2018; Jin *et al*., 2024), our infection assays demonstrate that its absence does not compromise TR4 virulence in banana. These findings indicate that FA’s role is context-dependent, potentially more significant in competitive microbial environments than in direct pathogenesis. Such flexibility highlights the adaptive resilience of TR4, which potentially compensates for FA loss through other genetic and metabolic pathways, for instance increased production of other antimicrobial compounds such as beauvericin, bikaverin, and equisetin to maintain ecological fitness (Spraker *et al*., 2018; Xu *et al*., 2023). Intriguingly, we also identified the loss of the FA BGC in another four *Fusarium* strains infecting common bean, suggesting that this cluster may be lost more frequently through diverse evolutionary mechanisms. Additionally, FA production has been shown to be dispensable for virulence in other pathosystems, such as the interaction between *F. verticillioides* and maize (Brown *et al*., 2015). In general, the levels of FA production between different *Fusarium* strains can differ significantly, irrespective of virulence against a host (Notz *et al*., 2002; Schouten *et al*., 2004). Together, these findings indicate that FA’s role in virulence is context-dependent, with its importance varying based on the host-pathogen system and environmental pressure.

Genomic variation because of TE activity, such as FoHeli1, may facilitate adaptation to environmental pressures, including host resistance and agricultural practices (Seidl & Thomma, 2017). This capacity for genomic remodeling, while beneficial for the pathogen, poses challenges for disease management, as it can lead to the emergence of new virulence traits or resistance mechanisms (Chuma *et al*., 2011; Kashiwa *et al*., 2016; Biju *et al*., 2017; Zaccaron & Stergiopoulos, 2024). At the same time unrestricted TE activity and proliferation might have long term consequences for the pathogen (Fouché *et al*., 2022). Expanding long-read genomic studies across the FOSC will allow for a better assessment of the impact of TEs on structural variation, gene loss and genome evolution.

Furthermore, the loss of FA production raises intriguing questions about the ecological trade-offs faced by TR4. The antimicrobial properties of FA likely confer advantages in niche competition (Notz *et al*., 2002; Bacon *et al*., 2006; Tung *et al*., 2017; Jin *et al*., 2024), yet its self-toxicity and energetic cost may render its production dispensable under certain conditions. The toxicity of FA has been linked to its chelating capacity, modulating the availability of metal ions essential for microbial survival (Ruiz *et al*., 2015; López-Díaz *et al*., 2018). On the other hand, other microbes have developed several ways to withstand higher concentrations of FA, such as efflux transporters or the increased production of siderophores (Hu *et al*., 2012; Ruiz *et al*., 2015). Studies showing reduced phytotoxic and antimicrobial effects of FA in the presence of metals like copper, iron and zinc support this hypothesis (Ruiz *et al*., 2015; López-Díaz *et al*., 2018). This trade-off mirrors the broader evolutionary dynamics of secondary metabolites in fungal pathogens, where ecological versatility often necessitates balancing metabolic costs and adaptive benefits (Brakhage, 2013; Krishnan *et al*., 2018; Abraham *et al*., 2024).

In conclusion, this study highlights the dual role of TEs in generating novel genetic variation and affecting genomic instability. In addition, we show that the conserved FA-BGC can be lost in TR4 without affecting virulence on banana. The ongoing activity of FoHeli1 in TR4 exemplifies the dynamic interplay between TE activity, genome evolution and ecological adaptation in fungal pathogens. By elucidating the consequences of TE-mediated genomic changes, we advance our understanding of the impact of TEs on genome dynamics and secondary metabolism in a clonally reproducing pathogen.

## Acknowledgements

JD, ACW, CAG, GNT and GHJK were supported by the Bill and Melinda Gates Foundation, grant no.: AG – 4425. Banana research at Wageningen University has been supported by the Dutch Dioraphte Foundation, grant no.: 20 04 04 02.

## Author contributions

JD conducted experiments and performed analyses. YL, CAG, and GNT conducted experiments. ACW, XSK and DWE contributed to the analyses. JD wrote the manuscript with input from GHJK and DWE. All authors contributed to writing and editing the manuscript. GHJK conceived and supervised the project.

## Data availability

All data are available in the main text or in the supporting information (Figs S1-S10; Tables S1-S4).

## References

Abraham LN, Oggenfuss U, Croll D. 2024. Population-level transposable element expression dynamics influence trait evolution in a fungal crop pathogen. mBio 15(3): e0284023.

Ayukawa Y, Asai S, Gan P, Tsushima A, Ichihashi Y, Shibata A, Komatsu K, Houterman PM, Rep M, Shirasu K, et al. 2021. A pair of effectors encoded on a conditionally dispensable chromosome of *Fusarium oxysporum* suppress host-specific immunity. Commun Biol 4(1): 707.

Bacon CW, Hinton DM, Hinton A, Jr. 2006. Growth-inhibiting effects of concentrations of fusaric acid on the growth of *Bacillus mojavensis* and other biocontrol *Bacillus* species. J Appl Microbiol 100(1): 185–194.

Baril T, Galbraith J, Hayward A. 2024. Earl Grey: A Fully Automated User-Friendly Transposable Element Annotation and Analysis Pipeline. Mol Biol Evol 41(4).

Barro-Trastoy D, Köhler C. 2024. Helitrons: genomic parasites that generate developmental novelties. Trends Genet 40(5): 437–448.

Bertazzoni S, Williams AH, Jones DA, Syme RA, Tan KC, Hane JK. 2018. Accessories Make the Outfit: Accessory Chromosomes and Other Dispensable DNA Regions in Plant-Pathogenic Fungi. Mol Plant Microbe Interact 31(8): 779–788.

Biju VC, Fokkens L, Houterman PM, Rep M, Cornelissen BJC. 2017. Multiple Evolutionary Trajectories Have Led to the Emergence of Races in *Fusarium oxysporum* f. sp. *lycopersici*. Appl Environ Microbiol 83(4).

Blin K, Shaw S, Augustijn HE, Reitz ZL, Biermann F, Alanjary M, Fetter A, Terlouw BR, Metcalf WW, Helfrich EJN, et al. 2023. antiSMASH 7.0: new and improved predictions for detection, regulation, chemical structures and visualisation. Nucleic Acids Res 51(W1): W46–W50.

Bohni N, Hofstetter V, Gindro K, Buyck B, Schumpp O, Bertrand S, Monod M, Wolfender JL. 2016. Production of Fusaric Acid by *Fusarium* spp. in Pure Culture and in Solid Medium Co-Cultures. Molecules 21(3): 370.

Brakhage AA. 2013. Regulation of fungal secondary metabolism. Nat Rev Microbiol 11(1): 21–32.

Brown DW, Lee SH, Kim LH, Ryu JG, Lee S, Seo Y, Kim YH, Busman M, Yun SH, Proctor RH, et al. 2015. Identification of a 12-gene Fusaric Acid Biosynthetic Gene Cluster in *Fusarium* Species Through Comparative and Functional Genomics. Mol Plant Microbe Interact 28(3): 319–332.

Camacho C, Coulouris G, Avagyan V, Ma N, Papadopoulos J, Bealer K, Madden TL. 2009. BLAST+: architecture and applications. BMC Bioinformatics 10: 421.

Castanera R, Lopez-Varas L, Borgognone A, LaButti K, Lapidus A, Schmutz J, Grimwood J, Perez G, Pisabarro AG, Grigoriev IV, et al. 2016. Transposable Elements versus the Fungal Genome: Impact on Whole-Genome Architecture and Transcriptional Profiles. PLoS Genet 12(6): e1006108.

Castanera R, Perez G, Lopez L, Sancho R, Santoyo F, Alfaro M, Gabaldon T, Pisabarro AG, Oguiza JA, Ramirez L. 2014. Highly expressed captured genes and cross-kingdom domains present in Helitrons create novel diversity in *Pleurotus ostreatus* and other fungi. BMC Genomics 15(1): 1071.

Catlin NS, Josephs EB. 2022. The important contribution of transposable elements to phenotypic variation and evolution. Curr Opin Plant Biol 65: 102140.

Chalopin D, Naville M, Plard F, Galiana D, Volff JN. 2015. Comparative analysis of transposable elements highlights mobilome diversity and evolution in vertebrates. Genome Biol Evol 7(2): 567–580.

Chao J, Li Z, Sun Y, Aluko OO, Wu X, Wang Q, Liu G. 2021. MG2C: a user-friendly online tool for drawing genetic maps. Mol Hortic 1(1): 16.

Chellapan BV, van Dam P, Rep M, Cornelissen BJ, Fokkens L. 2016. Non-canonical Helitrons in *Fusarium oxysporum*. Mob DNA 7: 27.

Chuma I, Isobe C, Hotta Y, Ibaragi K, Futamata N, Kusaba M, Yoshida K, Terauchi R, Fujita Y, Nakayashiki H, et al. 2011. Multiple translocation of the AVR-Pita effector gene among chromosomes of the rice blast fungus *Magnaporthe oryzae* and related species. PLoS Pathog 7(7): e1002147.

Crutcher FK, Liu J, Puckhaber LS, Stipanovic RD, Bell AA, Nichols RL. 2015. FUBT, a putative MFS transporter, promotes secretion of fusaric acid in the cotton pathogen *Fusarium oxysporum* f. sp. *vasinfectum*. Microbiology (Reading) 161(Pt 4): 875–883.

de Jonge R, Bolton MD, Kombrink A, van den Berg GC, Yadeta KA, Thomma BP. 2013. Extensive chromosomal reshuffling drives evolution of virulence in an asexual pathogen. Genome Res 23(8): 1271–1282.

De Vos RC, Moco S, Lommen A, Keurentjes JJ, Bino RJ, Hall RD. 2007. Untargeted large-scale plant metabolomics using liquid chromatography coupled to mass spectrometry. Nat Protoc 2(4): 778–791.

Dean R, Van Kan JA, Pretorius ZA, Hammond-Kosack KE, Di Pietro A, Spanu PD, Rudd JJ, Dickman M, Kahmann R, Ellis J, et al. 2012. The Top 10 fungal pathogens in molecular plant pathology. Mol Plant Pathol 13(4): 414–430.

Dijkstra J, van Westerhoven AC, Gómez-Gil L, Aguilera-Galvez C, Nakasato-Tagami G, Garnier SD, Yamazaki M, Arie T, Kamakura T, Arazoe T, et al. 2024. Extensive intrachromosomal duplications in a virulence-associated fungal accessory chromosome. bioRxiv: 2024.2009.2016.611982.

Du C, Fefelova N, Caronna J, He L, Dooner HK. 2009. The polychromatic Helitron landscape of the maize genome. Proc Natl Acad Sci U S A 106(47): 19916–19921.

Dührkop K, Fleischauer M, Ludwig M, Aksenov AA, Melnik AV, Meusel M, Dorrestein PC, Rousu J, Bocker S. 2019. SIRIUS 4: a rapid tool for turning tandem mass spectra into metabolite structure information. Nat Methods 16(4): 299–302.

Faino L, Seidl MF, Shi-Kunne X, Pauper M, van den Berg GC, Wittenberg AH, Thomma BP. 2016. Transposons passively and actively contribute to evolution of the two-speed genome of a fungal pathogen. Genome Res 26(8): 1091–1100.

Fouché S, Badet T, Oggenfuss U, Plissonneau C, Francisco CS, Croll D. 2020. Stress-Driven Transposable Element De-repression Dynamics and Virulence Evolution in a Fungal Pathogen. Mol Biol Evol 37(1): 221–239.

Fouché S, Oggenfuss U, Chanclud E, Croll D. 2022. A devil’s bargain with transposable elements in plant pathogens. Trends Genet 38(3): 222–230.

Frison EA, Escalant, J. V., Sharrock, S., Jain, S. M., & Swennen, R. . 2004. The global Musa genomic consortium: a boost for banana improvement.

García-Bastidas FA, Van der Veen AJT, Nakasato-Tagami G, Meijer HJG, Arango-Isaza RE, Kema GHJ. 2019. An Improved Phenotyping Protocol for Panama Disease in Banana. Front Plant Sci 10: 1006.

Gaumann E. 1957. Fusaric acid as a wilt toxin. Phytopathology 47(6): 342–357 pp.

Gilchrist CLM, Chooi YH. 2021. clinker & clustermap.js: automatic generation of gene cluster comparison figures. Bioinformatics 37(16): 2473–2475.

Gluck-Thaler E, Ralston T, Konkel Z, Ocampos CG, Ganeshan VD, Dorrance AE, Niblack TL, Wood CW, Slot JC, Lopez-Nicora HD, et al. 2022. Giant Starship Elements Mobilize Accessory Genes in Fungal Genomes. Mol Biol Evol 39(5).

Grabundzija I, Messing SA, Thomas J, Cosby RL, Bilic I, Miskey C, Gogol-Doring A, Kapitonov V, Diem T, Dalda A, et al. 2016. A Helitron transposon reconstructed from bats reveals a novel mechanism of genome shuffling in eukaryotes. Nat Commun 7: 10716.

Hayward A, Gilbert C. 2022. Transposable elements. Curr Biol 32(17): R904–R909.

Hirsch CD, Springer NM. 2017. Transposable element influences on gene expression in plants. Biochim Biophys Acta Gene Regul Mech 1860(1): 157–165.

Hu K, Xu K, Wen J, Yi B, Shen J, Ma C, Fu T, Ouyang Y, Tu J. 2019. Helitron distribution in *Brassicaceae* and whole Genome Helitron density as a character for distinguishing plant species. BMC Bioinformatics 20(1): 354.

Hu RM, Liao ST, Huang CC, Huang YW, Yang TC. 2012. An inducible fusaric acid tripartite efflux pump contributes to the fusaric acid resistance in *Stenotrophomonas maltophilia*. PLoS One 7(12): e51053.

Jin X, Jia H, Ran L, Wu F, Liu J, Schlaeppi K, Dini-Andreote F, Wei Z, Zhou X. 2024. Fusaric acid mediates the assembly of disease-suppressive rhizosphere microbiota via induced shifts in plant root exudates. Nat Commun 15(1): 5125.

Karamura E, Frison, E., Karamura, D. A., & Sharrock, S.. 1998. Banana production systems in eastern and southern Africa.

Karangwa P, Blomme G, Beed F, Niyongere C, Viljoen A. 2016. The distribution and incidence of banana Fusarium wilt in subsistence farming systems in east and central Africa. Crop Protection 84: 132–140.

Kashiwa T, Suzuki T, Sato A, Akai K, Teraoka T, Komatsu K, Arie T. 2016. A new biotype of *Fusarium oxysporum* f. sp. *lycopersici* race 2 emerged by a transposon-driven mutation of avirulence gene *AVR1*. FEMS Microbiol Lett 363(14).

Krishnan P, Meile L, Plissonneau C, Ma X, Hartmann FE, Croll D, McDonald BA, Sanchez-Vallet A. 2018. Transposable element insertions shape gene regulation and melanin production in a fungal pathogen of wheat. BMC Biol 16(1): 78.

Li C, Zuo C, Deng G, Kuang R, Yang Q, Hu C, Sheng O, Zhang S, Ma L, Wei Y, et al. 2013. Contamination of bananas with beauvericin and fusaric acid produced by *Fusarium oxysporum* f. sp. *cubense*. PLoS One 8(7): e70226.

Li J, Fokkens L, van Dam P, Rep M. 2020. Related mobile pathogenicity chromosomes in *Fusarium oxysporum* determine host range on cucurbits. Mol Plant Pathol 21(6): 761–776.

Li Y, Dooner HK. 2009. Excision of Helitron transposons in maize. Genetics 182(1): 399–402.

Li Z, Gilbert C, Peng H, Pollet N. 2024. Discovery of numerous novel Helitron-like elements in eukaryote genomes using HELIANO. Nucleic Acids Res 52(17): e79.

Liu S, Li J, Zhang Y, Liu N, Viljoen A, Mostert D, Zuo C, Hu C, Bi F, Gao H, et al. 2020. Fusaric acid instigates the invasion of banana by *Fusarium oxysporum* f. sp. *cubense* TR4. New Phytol 225(2): 913–929.

Lommen A. 2009. MetAlign: interface-driven, versatile metabolomics tool for hyphenated full-scan mass spectrometry data preprocessing. Anal Chem 81(8): 3079–3086.

López-Díaz C, Rahjoo V, Sulyok M, Ghionna V, Martin-Vicente A, Capilla J, Di Pietro A, Lopez-Berges MS. 2018. Fusaric acid contributes to virulence of *Fusarium oxysporum* on plant and mammalian hosts. Mol Plant Pathol 19(2): 440–453.

Ma LJ, van der Does HC, Borkovich KA, Coleman JJ, Daboussi MJ, Di Pietro A, Dufresne M, Freitag M, Grabherr M, Henrissat B, et al. 2010. Comparative genomics reveals mobile pathogenicity chromosomes in *Fusarium*. Nature 464(7287): 367–373.

Martínez-de la Parte E, Perez-Vicente L, Torres DE, van Westerhoven A, Meijer HJG, Seidl MF, Kema GHJ. 2024. Genetic diversity of the banana Fusarium wilt pathogen in Cuba and across Latin America and the Caribbean. Environ Microbiol 26(5): e16636.

Meng T, Jiao H, Zhang Y, Zhou Y, Chen S, Wang X, Yang B, Sun J, Geng X, Ayhan DH, et al. 2024. FoPGDB: a pangenome database of *Fusarium oxysporum*, a cross-kingdom fungal pathogen. Database (Oxford) 2024.

Möller M, Stukenbrock EH. 2017. Evolution and genome architecture in fungal plant pathogens. Nat Rev Microbiol 15(12): 756–771.

Niehaus EM, von Bargen KW, Espino JJ, Pfannmuller A, Humpf HU, Tudzynski B. 2014. Characterization of the fusaric acid gene cluster in *Fusarium fujikuroi*. Appl Microbiol Biotechnol 98(4): 1749–1762.

Notz R, Maurhofer M, Dubach H, Haas D, Defago G. 2002. Fusaric acid-producing strains of *Fusarium oxysporum* alter 2,4-diacetylphloroglucinol biosynthetic gene expression in *Pseudomonas fluorescens* CHA0 in vitro and in the rhizosphere of wheat. Appl Environ Microbiol 68(5): 2229–2235.

Oggenfuss U, Badet T, Croll D. 2024. A systematic screen for co-option of transposable elements across the fungal kingdom. Mob DNA 15(1): 2.

Ordóñez N, Seidl MF, Waalwijk C, Drenth A, Kilian A, Thomma BP, Ploetz RC, Kema GH. 2015. Worse Comes to Worst: Bananas and Panama Disease--When Plant and Pathogen Clones Meet. PLoS Pathog 11(11): e1005197.

Phasha MM, Wingfield BD, Wingfield MJ, Coetzee MPA, Hammerbacher A, Steenkamp ET. 2021. Deciphering the effect of FUB1 disruption on fusaric acid production and pathogenicity in *Fusarium circinatum*. Fungal Biol 125(12): 1036–1047.

Ploetz RC. 2015. Management of Fusarium wilt of banana: A review with special reference to tropical race 4. Crop Protection 73: 7–15.

Ploetz RC, Kema GHJ, Ma L-J. 2015. Impact of Diseases on Export and Smallholder Production of Banana. Annual Review of Phytopathology 53(1): 269–288.

Quigley S, Damas J, Larkin DM, Farre M. 2023. syntenyPlotteR: a user-friendly R package to visualize genome synteny, ideal for both experienced and novice bioinformaticians. Bioinform Adv 3(1): vbad161.

Rouxel T, Grandaubert J, Hane JK, Hoede C, van de Wouw AP, Couloux A, Dominguez V, Anthouard V, Bally P, Bourras S, et al. 2011. Effector diversification within compartments of the *Leptosphaeria maculan*s genome affected by Repeat-Induced Point mutations. Nat Commun 2: 202.

Ruiz JA, Bernar EM, Jung K. 2015. Production of siderophores increases resistance to fusaric acid in *Pseudomonas protegens* Pf-5. PLoS One 10(1): e0117040.

Schmid R, Heuckeroth S, Korf A, Smirnov A, Myers O, Dyrlund TS, Bushuiev R, Murray KJ, Hoffmann N, Lu M, et al. 2023. Integrative analysis of multimodal mass spectrometry data in MZmine 3. Nat Biotechnol 41(4): 447–449.

Schmidt SM, Houterman PM, Schreiver I, Ma L, Amyotte S, Chellappan B, Boeren S, Takken FL, Rep M. 2013. MITEs in the promoters of effector genes allow prediction of novel virulence genes in *Fusarium oxysporum*. BMC Genomics 14: 119.

Schouten A, van den Berg G, Edel-Hermann V, Steinberg C, Gautheron N, Alabouvette C, de Vos CH, Lemanceau P, Raaijmakers JM. 2004. Defense responses of *Fusarium oxysporum* to 2,4-diacetylphloroglucinol, a broad-spectrum antibiotic produced by *Pseudomonas fluorescens*. Mol Plant Microbe Interact 17(11): 1201–1211.

Seidl MF, Thomma B. 2017. Transposable Elements Direct The Coevolution between Plants and Microbes. Trends Genet 33(11): 842–851.

Sievers F, Wilm A, Dineen D, Gibson TJ, Karplus K, Li W, Lopez R, McWilliam H, Remmert M, Söding J, et al. 2011. Fast, scalable generation of high-quality protein multiple sequence alignments using Clustal Omega. Mol Syst Biol 7: 539.

Son SW, Kim HY, Choi GJ, Lim HK, Jang KS, Lee SO, Lee S, Sung ND, Kim JC. 2008. Bikaverin and fusaric acid from *Fusarium oxysporum* show antioomycete activity against *Phytophthora infestans*. J Appl Microbiol 104(3): 692–698.

Spraker JE, Wiemann P, Baccile JA, Venkatesh N, Schumacher J, Schroeder FC, Sanchez LM, Keller NP. 2018. Conserved Responses in a War of Small Molecules between a Plant-Pathogenic Bacterium and Fungi. mBio 9(3).

Studt L, Janevska S, Niehaus EM, Burkhardt I, Arndt B, Sieber CM, Humpf HU, Dickschat JS, Tudzynski B. 2016. Two separate key enzymes and two pathway-specific transcription factors are involved in fusaric acid biosynthesis in *Fusarium fujikuroi*. Environ Microbiol 18(3): 936–956.

Thomas J, Pritham EJ. 2015. Helitrons, the Eukaryotic Rolling-circle Transposable Elements. Microbiol Spectr 3(4).

Tikunov YM, Laptenok S, Hall RD, Bovy A, de Vos RC. 2012. MSClust: a tool for unsupervised mass spectra extraction of chromatography-mass spectrometry ion-wise aligned data. Metabolomics 8(4): 714–718.

Torres DE, Oggenfuss U, Croll D, Seidl MF. 2020. Genome evolution in fungal plant pathogens: looking beyond the two-speed genome model. Fungal Biology Reviews 34(3): 136–143.

Tung TT, Jakobsen TH, Dao TT, Fuglsang AT, Givskov M, Christensen SB, Nielsen J. 2017. Fusaric acid and analogues as Gram-negative bacterial quorum sensing inhibitors. Eur J Med Chem 126: 1011–1020.

van Westerhoven AC, Aguilera-Galvez C, Nakasato-Tagami G, Shi-Kunne X, Martinez de la Parte E, Chavarro-Carrero E, Meijer HJG, Feurtey A, Maryani N, Ordóñez N, et al. 2024. Segmental duplications drive the evolution of accessory regions in a major crop pathogen. New Phytol 242(2): 610–625.

van Westerhoven AC, Meijer HJG, Houdijk J, Martinez de la Parte E, Matabuana EL, Seidl MF, Kema GHJ. 2023. Dissemination of Fusarium Wilt of Banana in Mozambique Caused by *Fusarium odoratissimum* Tropical Race 4. Plant Dis 107(3): 628–632.

van Westerhoven AC, Meijer HJG, Seidl MF, Kema GHJ. 2022. Uncontained spread of Fusarium wilt of banana threatens African food security. PLoS Pathog 18(9): e1010769.

Weilguny L, Kofler R. 2019. DeviaTE: Assembly-free analysis and visualization of mobile genetic element composition. Mol Ecol Resour 19(5): 1346–1354.

Wells JN, Feschotte C. 2020. A Field Guide to Eukaryotic Transposable Elements. Annu Rev Genet 54: 539–561.

Xu M, Huang Z, Zhu W, Liu Y, Bai X, Zhang H. 2023. *Fusarium*-Derived Secondary Metabolites with Antimicrobial Effects. Molecules 28(8).

Zaccaron AZ, Stergiopoulos I. 2024. Analysis of five near-complete genome assemblies of the tomato pathogen *Cladosporium fulvum* uncovers additional accessory chromosomes and structural variations induced by transposable elements effecting the loss of avirulence genes. BMC Biol 22(1): 25.

Zhang Y, Liu S, Mostert D, Yu H, Zhuo M, Li G, Zuo C, Haridas S, Webster K, Li M, et al. 2024. Virulence of banana wilt-causing fungal pathogen *Fusarium oxysporum* tropical race 4 is mediated by nitric oxide biosynthesis and accessory genes. Nat Microbiol.

