## Supplementary figures S1-10 for "Transposon activity eliminates a crucial fungal secondary metabolite cluster while preserving pathogenicity"

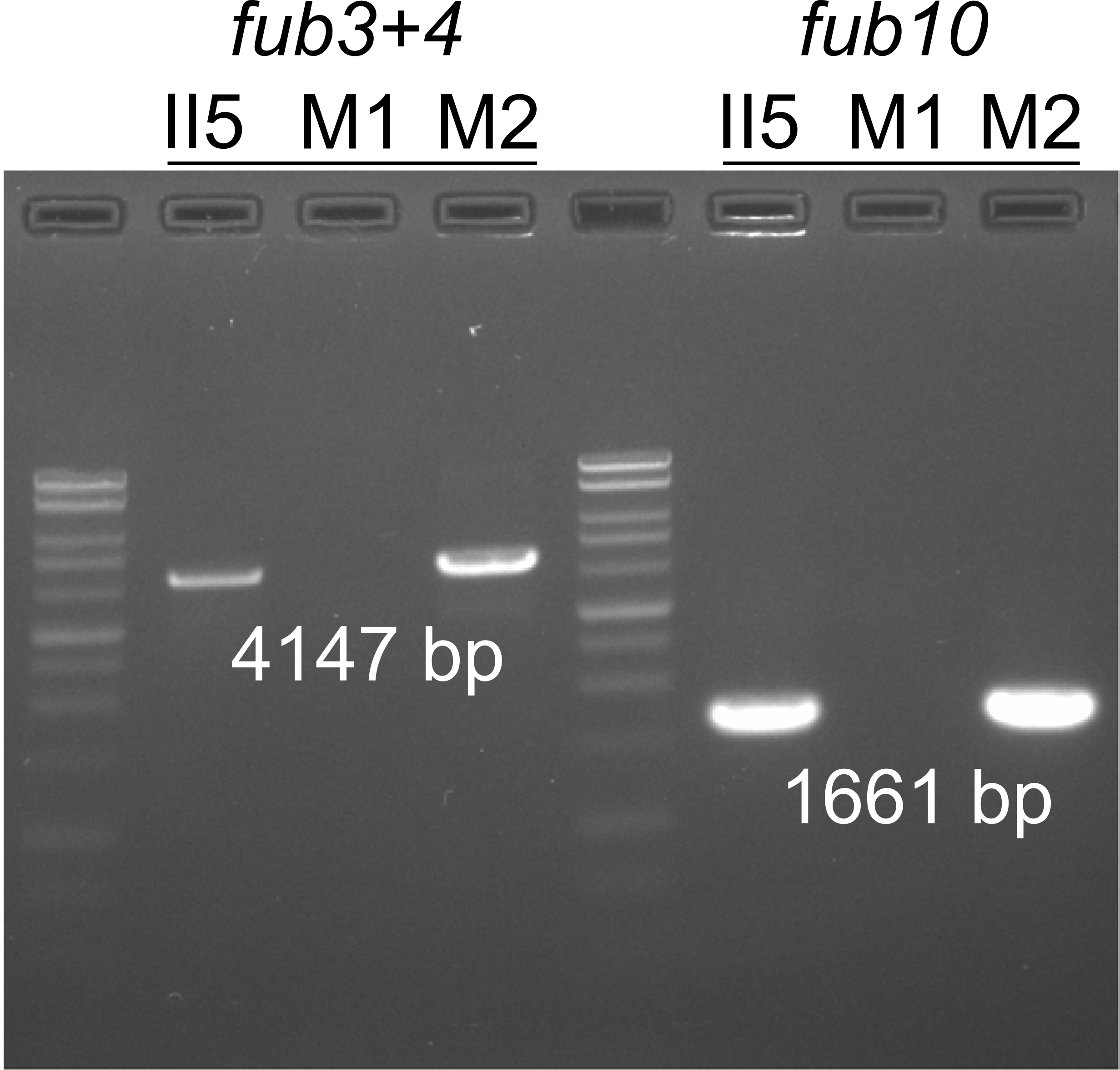

**Fig. S1. - PCR confirms loss of *fub* genes in TR4 strain M1.** Gel electrophoresis after PCR using primers for fragments containing *fub* genes 3+4 or *fub10* in TR4 strains II5, M1, and M2.

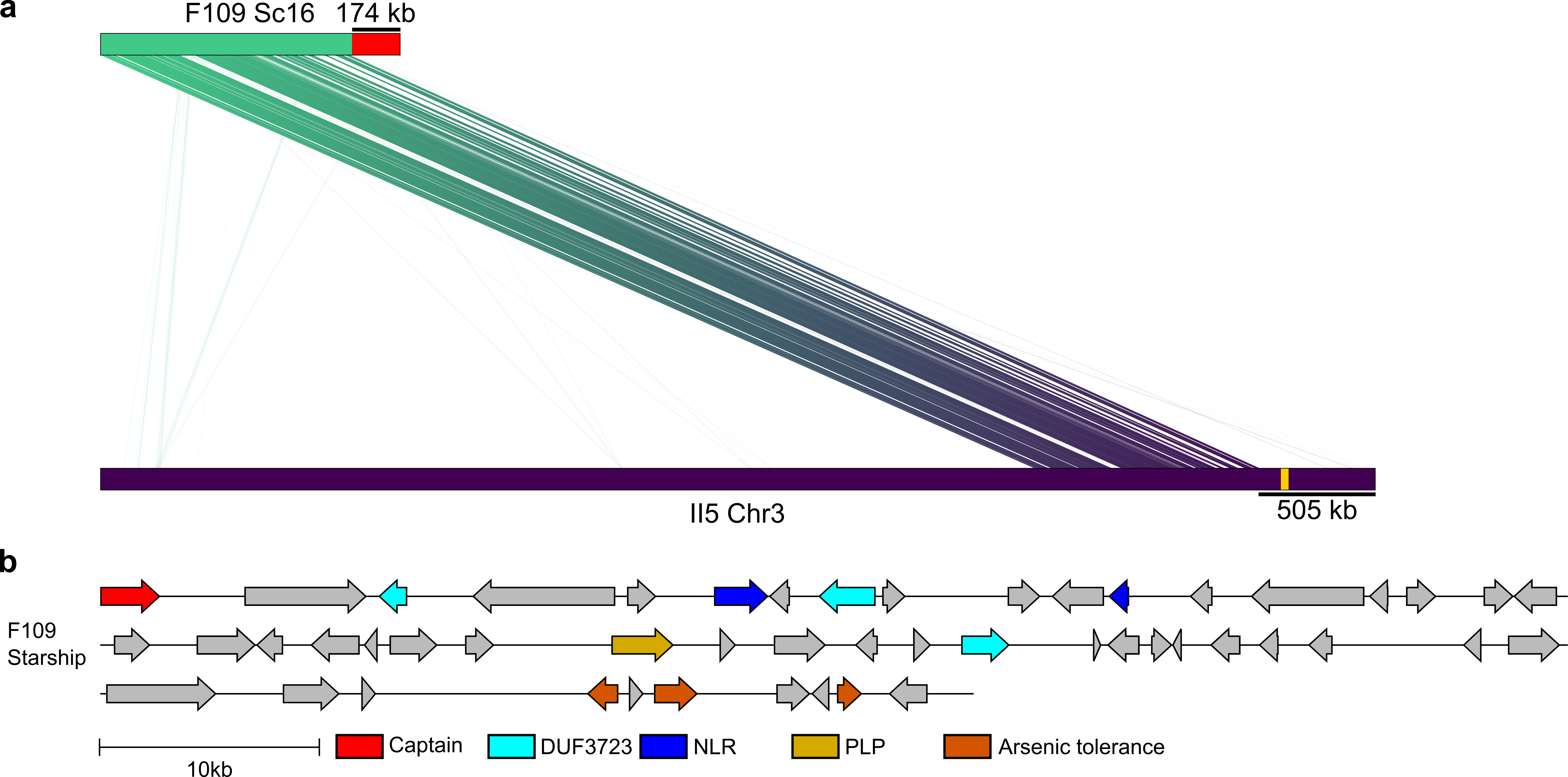
**Fig. S2. – Starship element in strain F109 caused fusaric acid BGC loss.** **a**) Alignment between TR4 strain II5 chromosome 3 and F109 strain scaffold 16. The approximate length of absent sequences is shown. The approximate location of the fusaric acid biosynthetic gene cluster is noted on II5 chromosome 3 by a yellow marker. The Starship element is depicted by a red marker on scaffold 16 of F109. The fusaric acid biosynthetic gene cluster is similarly absent in F23, F79 and F105 (Meng et al., 2024). **b**) Schematic representation of the 50 putative genes located on the Starship element on scaffold 16 of F109. Common Starship genes are noted (Captain=DUF3435, NLR=NOD-like receptor, PLP=patatin-like phospholipase). The scale bar depicts 10kb in length.

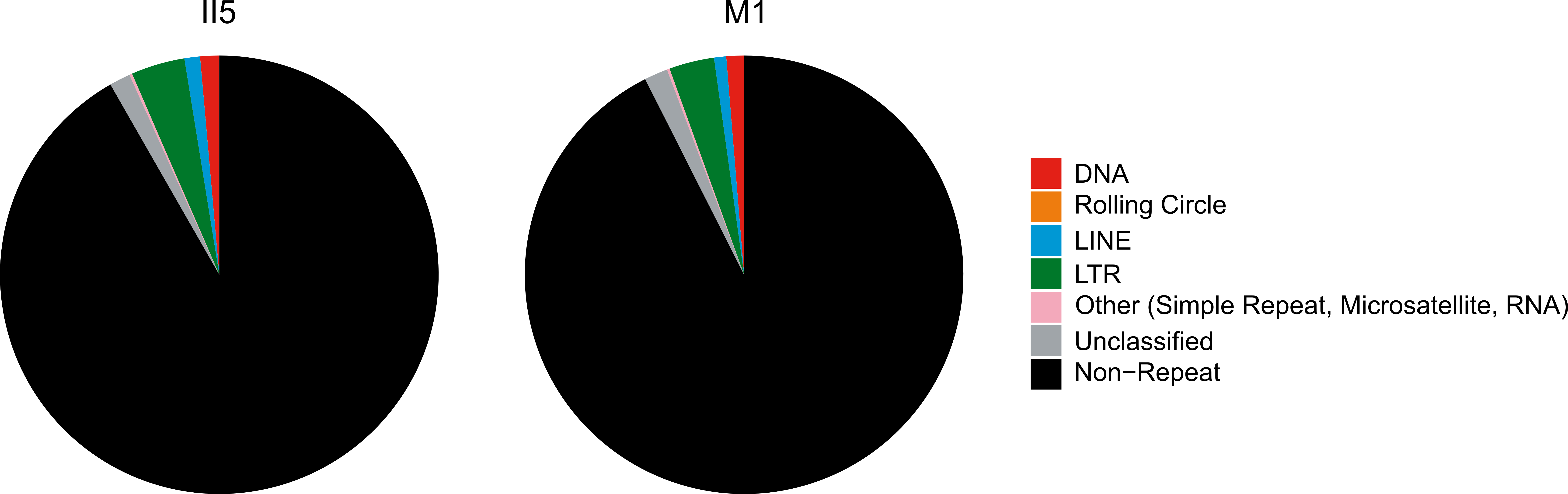
**Fig. S3. - Transposable element content of TR4 strains II5 and M1.** Pie chart summary of all TEs predicted by Earl Grey and their contribution to the total genome size of TR4 strains II5 (8.2%) and M1 (7.4%). Note FoHeli1 is part of the unclassified section.

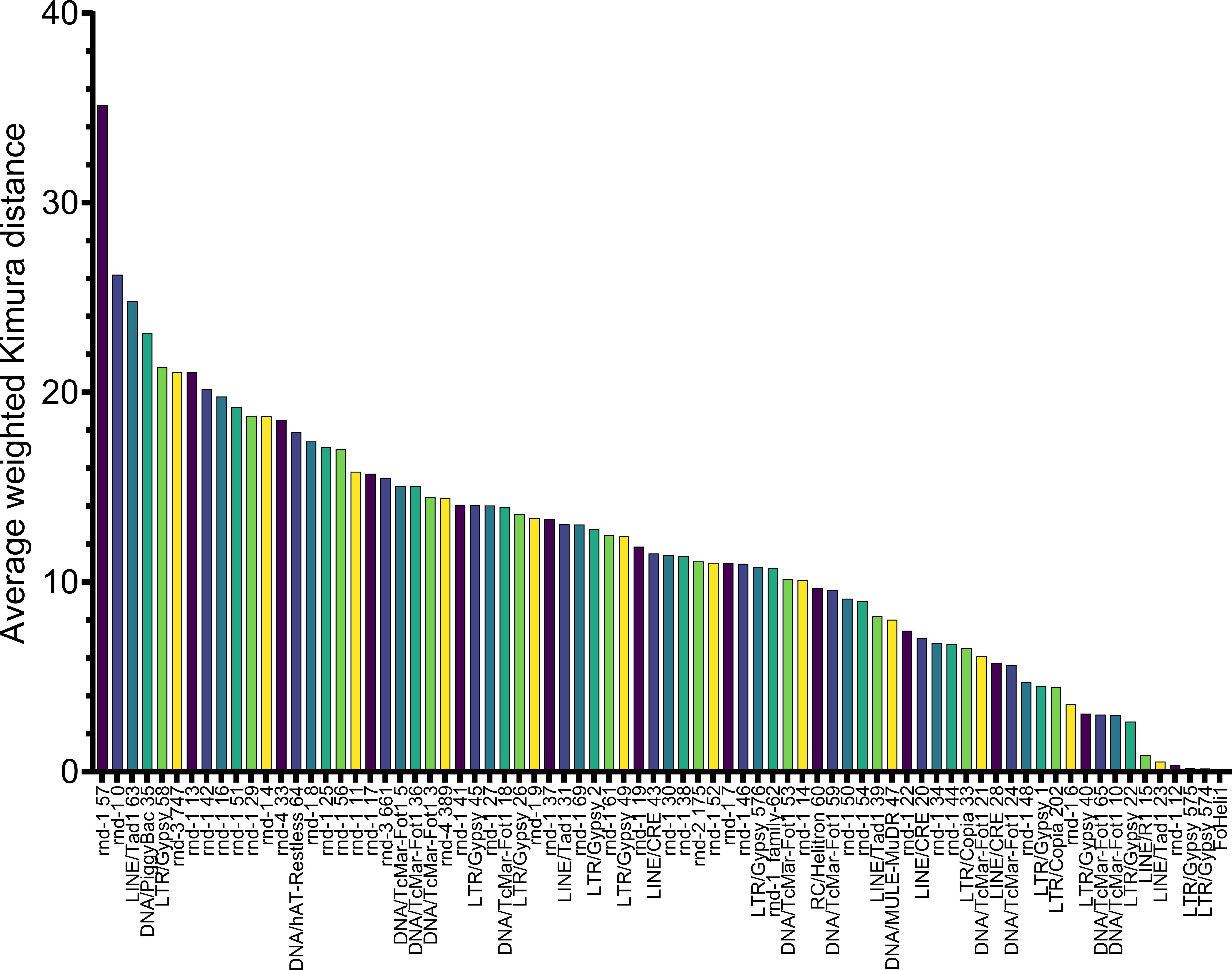

**Fig. S4. - Recent FoHeli1 activity in TR4 strain II5.** Bar chart showing the average weighted Kimura distance of all TEs predicted and calculated by Earl Grey in TR4 strain II5.

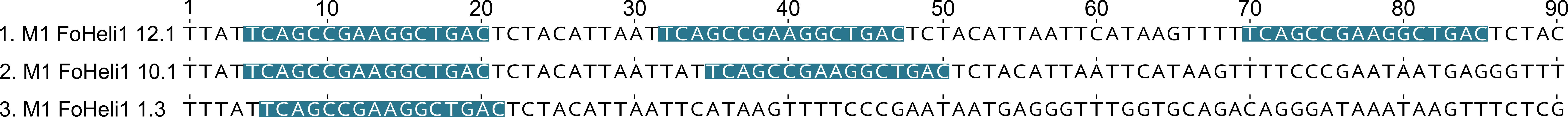
**Fig. S5. – M1 FoHeli1 with multiple 5’ termini.** First 100 bp of three FoHeli1 copies from TR4 strain M1 as predicted by Earl Grey. Colored markings indicate the 16 bp 5’ terminal sequence of FoHeli1 as described by Chellapan *et al*. (2016). FoHeli1 copies in M1 contain either one, two or three 5’ termini.

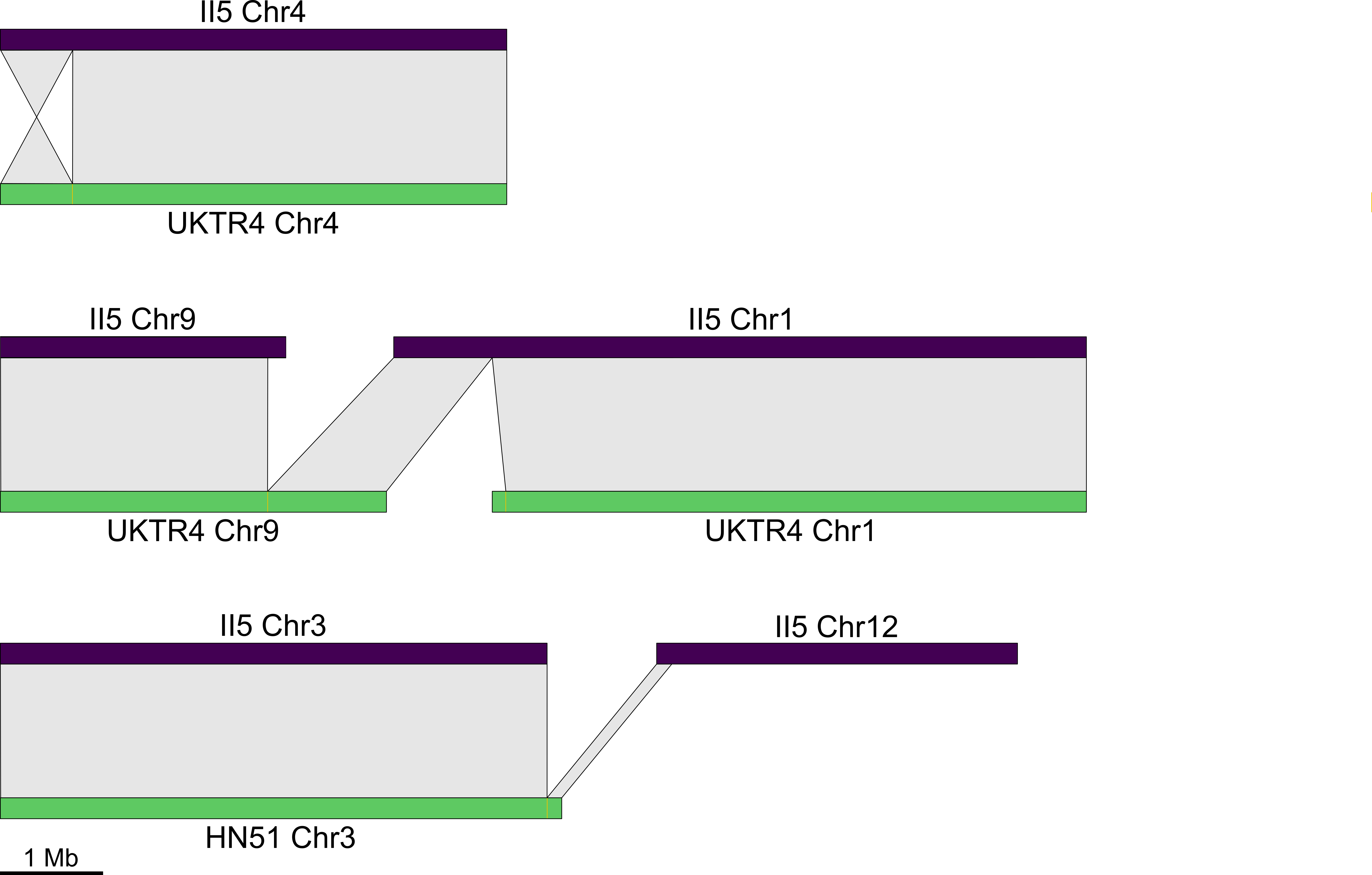
**Fig. S6. – FoHeli1 mediated genomic rearrangements in TR4 strains.** Alignment between chromosomes of TR4 strain II5, UKTR4 and HN51 showing rearrangements at FoHeli1 loci.

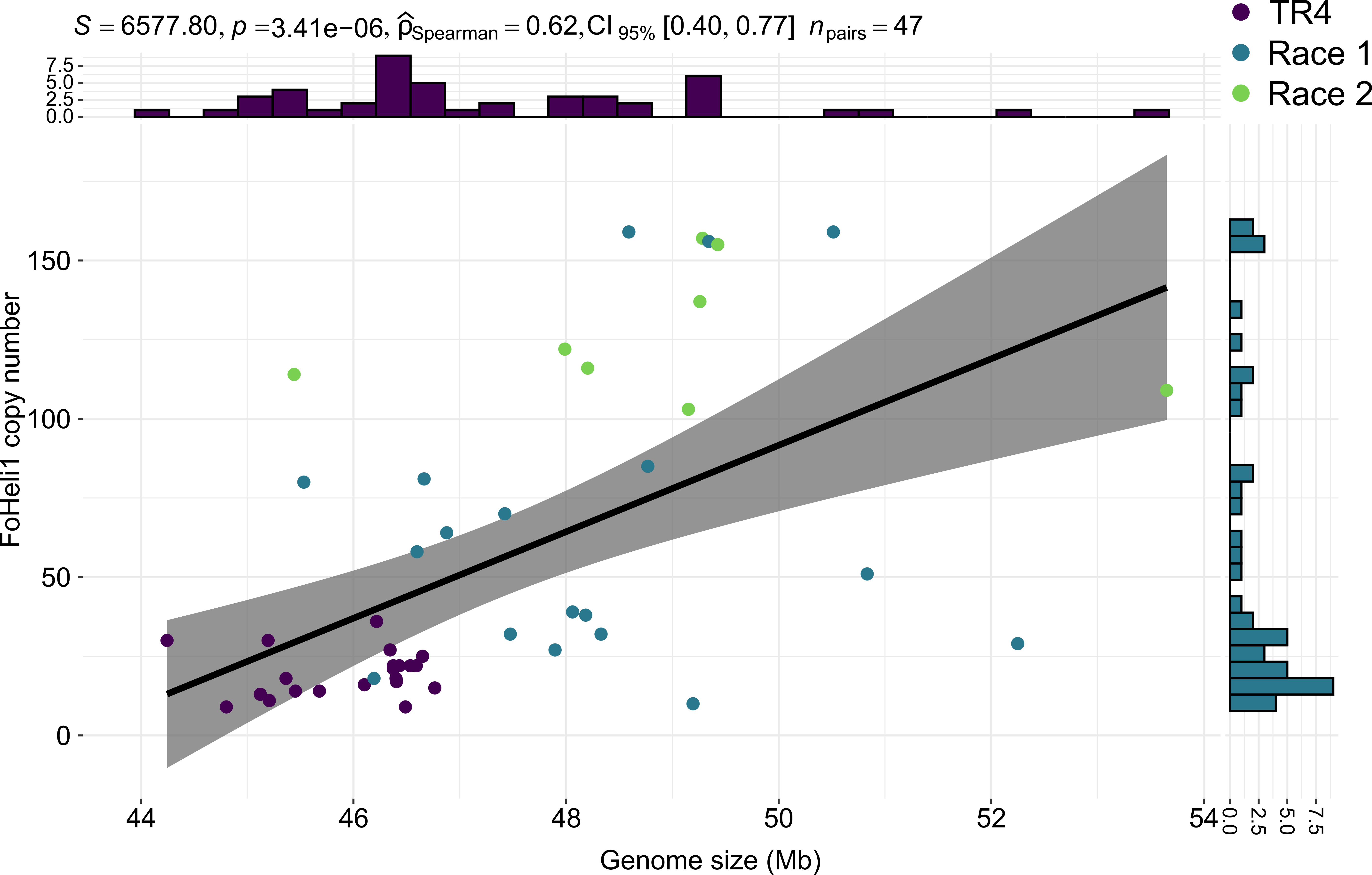

**Fig. S7. – Genome size shows moderate correlation with FoHeli1 copy number.** Comparison between genome size in Mbp and estimated FoHeli1 copy number for 47 banana infecting *Fusarium* strains containing at least one copy. Spearman correlation coefficient ($\hat{P}$), p value and 95% confidence interval are noted above.

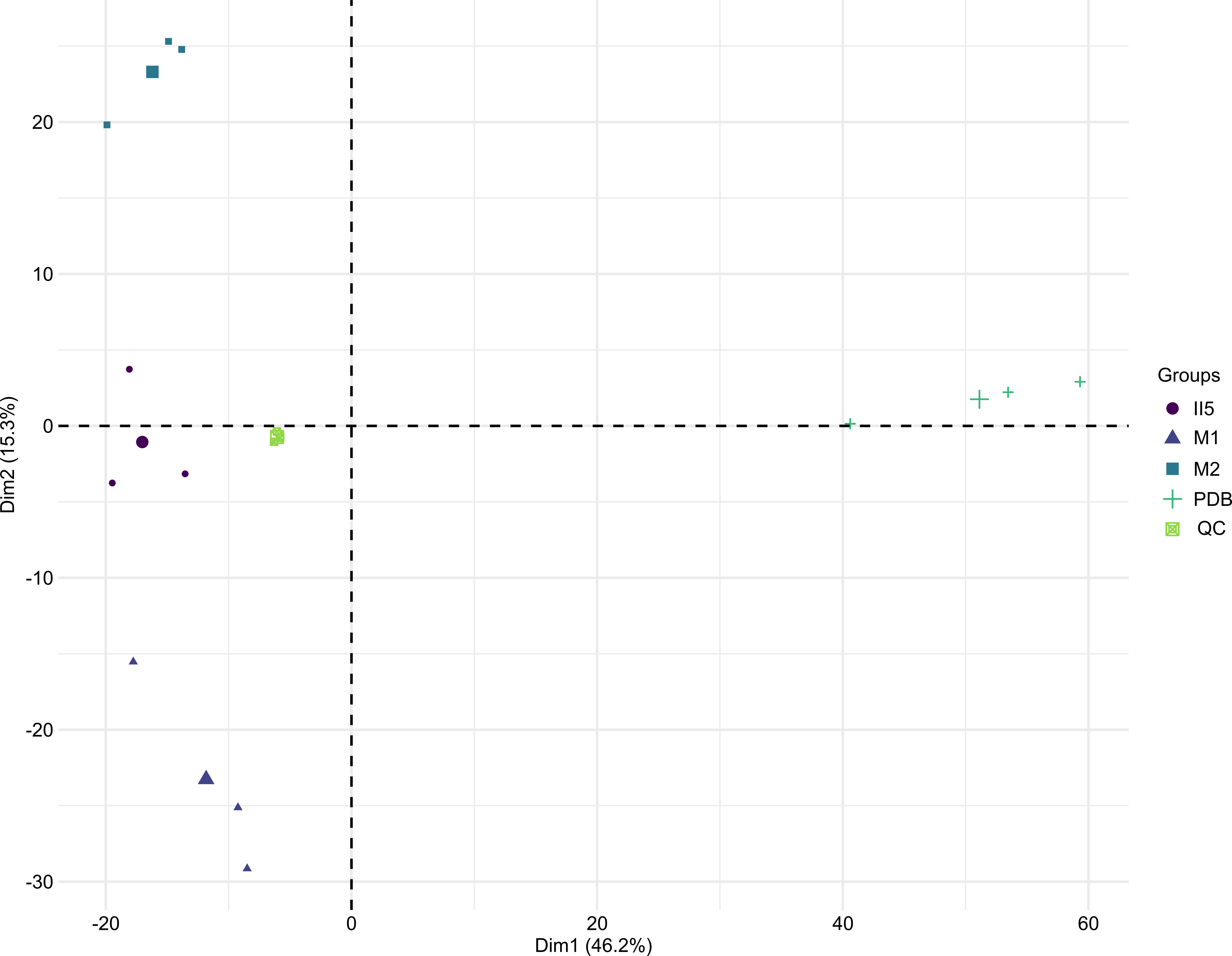

**Fig. S8. – TR4 strains II5, M1 and M2 show different exometabolite profiles.** PCA of the exometabolite profiles of TR4 strains II5, M1 and M2 as well as medium without fungal growth (PDB) and quality control samples (QC).

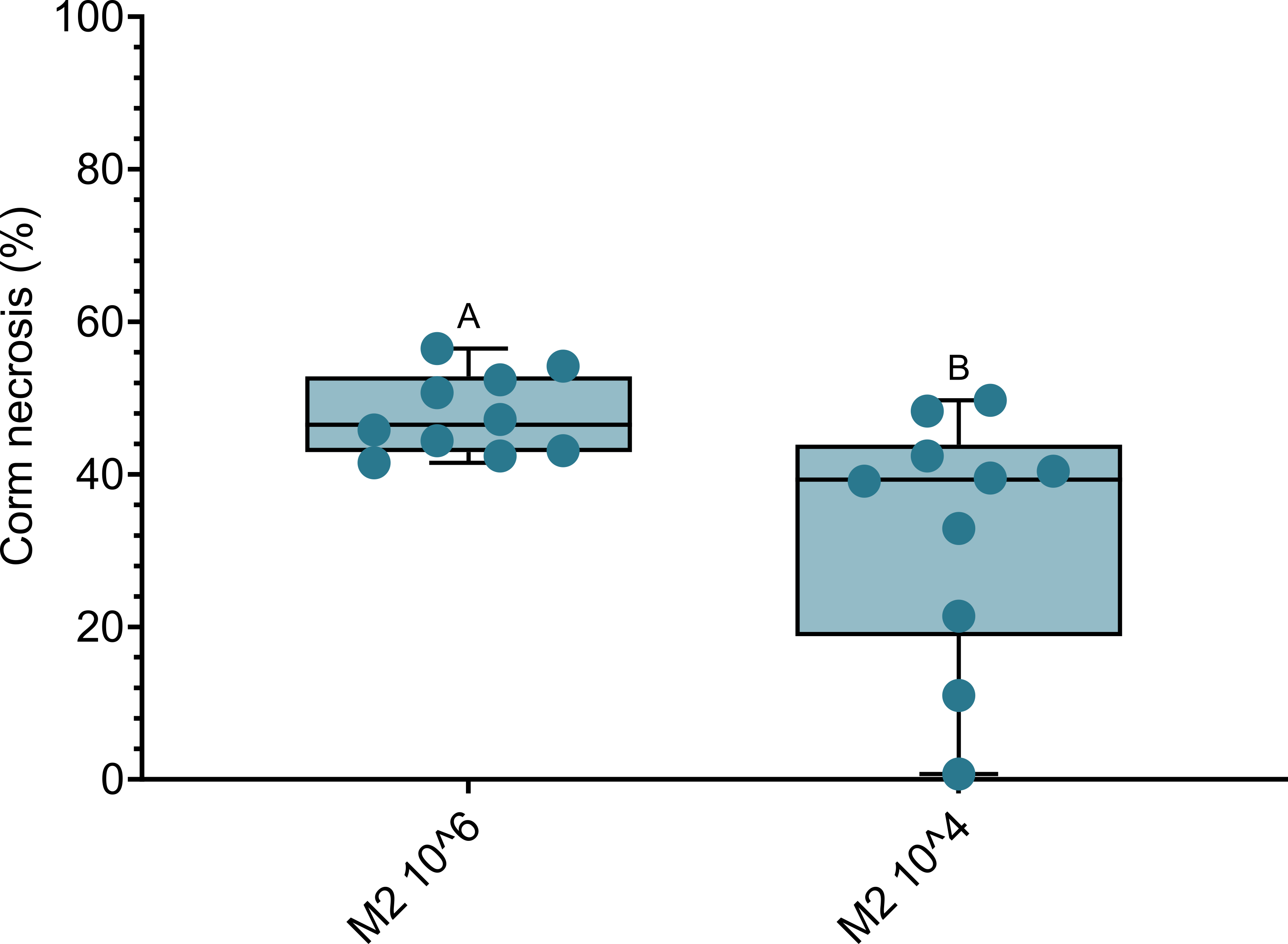

**Fig. S9. – M2 wilting severity is also affected by spore concentration.** Percentage of corm necrosis at 10 weeks post-inoculation of Cavendish 'Grand Naine' plants inoculated with TR4 strain M2. Plants were inoculated with either 10^6^ or 10^4^ spores per mL. Corm necrosis was quantified using ImageJ (n=10). Letters indicate significant differences between treatments (t-test; P<0.05).

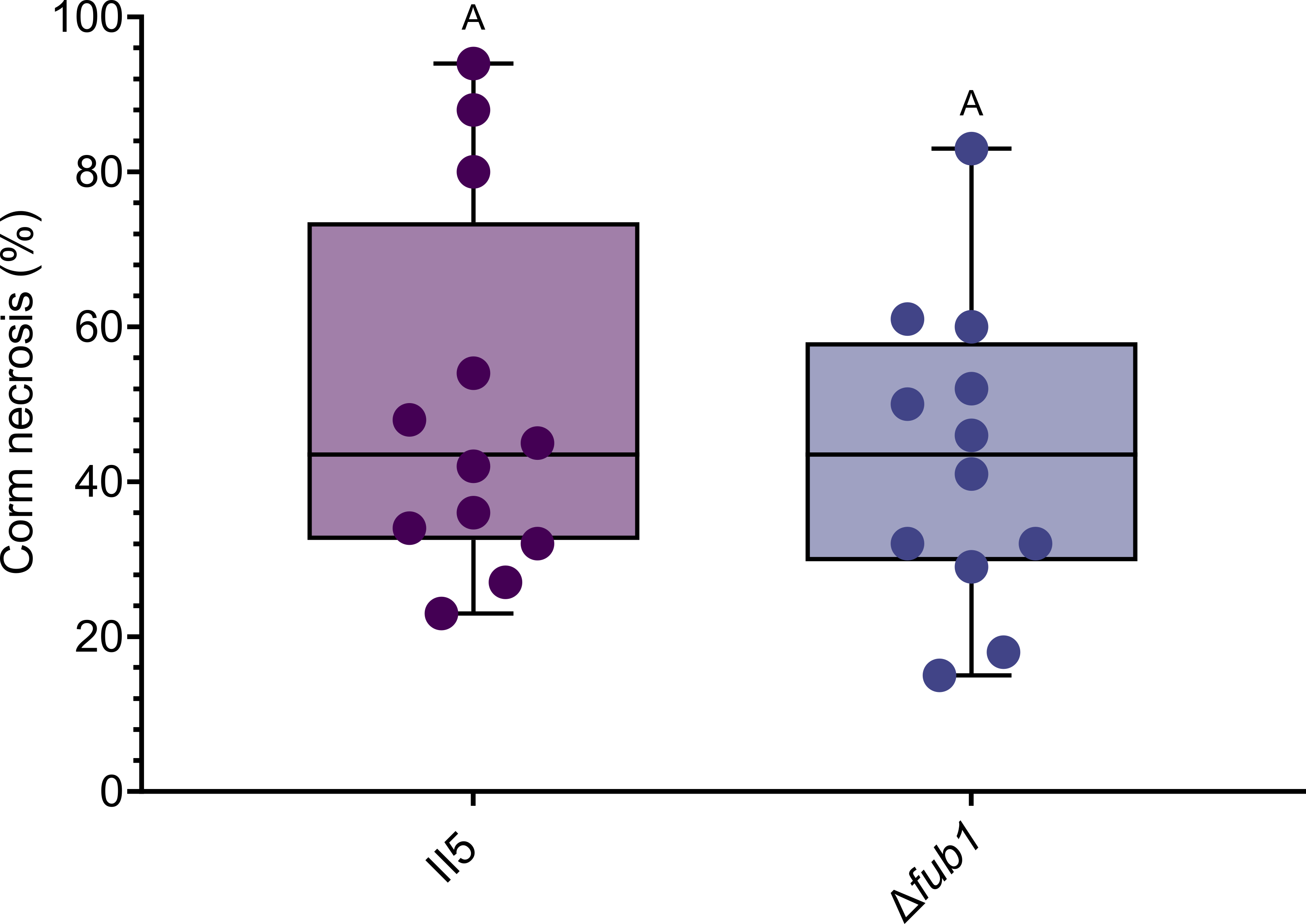

**Fig. S10. – Fusaric acid loss does not affect virulence at standard spore concentration.** Percentage of corm necrosis at 10 weeks post- inoculation of Cavendish 'Grand Naine' plants inoculated with reference strain II5 or Δ*fub1* mutant. All plants were inoculated with 10^6^ spores per mL. Corm necrosis was quantified using ImageJ (n=12). Letters indicate significant differences between treatments (t-test; P<0.05).
